# Towards Practical and Robust DNA-Based Data Archiving Using ‘Yin-Yang Codec’ System

**DOI:** 10.1101/829721

**Authors:** Zhi Ping, Shihong Chen, Guangyu Zhou, Xiaoluo Huang, Sha Joe Zhu, Haoling Zhang, Henry H. Lee, Zhaojun Lan, Jie Cui, Tai Chen, Wenwei Zhang, Huanming Yang, Xun Xu, George M. Church, Yue Shen

**Author notes:** These authors contributed equally to this work (Zhi Ping, Shihong Chen, Guangyu Zhou, Xiaoluo Huang).

## Abstract

DNA is a promising data storage medium due to its remarkable durability and space-efficient storage. Early bit-to-base transcoding schemes have primarily pursued information density, at the expense however of introducing biocompatibility challenges or at the risk of decoding failure. Here, we propose a robust transcoding algorithm named the “Yin-Yang Codec” (YYC), using two rules to encode two binary bits into one nucleotide, to generate DNA sequences highly compatible with synthesis and sequencing technologies. We encoded two representative file formats and stored them in vitro as 200-nt oligo pools and in vivo as an ~54-kb DNA fragment in yeast cells. Sequencing results show that YYC exhibits high robustness and reliability for a wide variety of data types, with an average recovery rate of 99.94% at 10^4^ molecule copies and an achieved recovery rate of 87.53% at 100 copies. In addition, the in vivo storage demonstration achieved for the first time an experimentally measured physical information density of 198.8 EB per gram of DNA (44% of the theoretical maximum for DNA).

## INTRODUCTION

DNA is an ancient and efficient information carrier in living organisms. At present, it is thought to have great potential as a novel storage medium, because standard storage media can no longer meet the exponentially increasing data archiving demands. The DNA molecule, compared to common information carriers, exhibits multiple advantages, including extremely high storage density (estimated physical density of 455 EB per gram of DNA (1)), extraordinary durability (half-life > 500 years (2,3)), and the capacity for cost-efficient information amplification.

Many strategies have been proposed for digital information storage using organic molecules, including DNA, oligopeptides, and metabolomes (4–9). Recently, DNA origami technology was also proposed as a novel strategy to store digital information using DNA nanostructures. Because current DNA sequencing technology holds advantages for both cost and throughput compared with mass spectroscopy or super-resolution microscopy technologies, storing digital information using DNA molecules remains the most well-accepted strategy. Binary information from each file is transcoded directly into DNA sequences, which are synthesized and stored in the form of oligonucleotides or double-stranded DNA fragments *in vitro* or *in vivo*. Then sequencing technology is used to retrieve stored digital information. In addition, several different molecular strategies have been proposed to implement the selective access of portions of the stored data to improve the practicality and scalability of DNA data storage (10–12).

Using basic transcoding rules (i.e., converting {00,01,10,11} to {A,C,G,T}), some specific patterns in DNA sequences will be generated that will introduce challenges to synthesis and sequencing (10,13,14). For example, single-nucleotide repeats (homopolymer) longer than 5-nt might introduce a higher error rate during sequencing or synthesis (15,16). It has also been reported that DNA sequences with stable secondary structure can create a disadvantage for sequencing to read the binary information or for PCR used for the random-access and backup of stored information (17–20). Additionally, DNA sequences with GC content < 40% or > 60% often create difficulties in DNA synthesis. Therefore, the length of homopolymers (in nt), the secondary structure (represented by free energy calculation, in kJ/mol), and the GC content (in %) are three primary parameters for evaluating the compatibility of coding schemes.

Previous studies on transcoding algorithm development attempted to improve the compatibility of generated DNA sequences. Early efforts, including those of Church *et al*. and Grass *et al*., introduced additional restrictions in the transcoding rules to eliminate homopolymers; however, this came at the expense of reduced information density (1,21,22). Later studies pioneered the base conversion rules without compromising information density. As a demonstration, DNA fountain adopted Luby Transform (LT) codes to improve the information fidelity by introducing low redundancy and screening constraints on the length of homopolymers and the GC content while maintaining an information density of 1.57 bits/nt (6,23). However, the major drawback for DNA fountain is that it risks unsuccessful decoding when dealing with particular binary features due to the fundamental issues of LT codes. Thus, it relies on the introduction of sufficient logical redundancy to ensure successful decoding. Logical redundancy refers to redundancy at the coding level, and this is different from physical redundancy, which refers to excessive synthesized DNA molecules. Reducing the logical redundancy could lead to a high probability of decoding failure, but excessive logical redundancy will decrease the information density and significantly increase the cost of synthesis (16). Furthermore, specific binary patterns using these early-established algorithms may also create unsuitable DNA sequences, either with extreme GC content or long homopolymers (Supplementary Table S1). Therefore, developing a coding algorithm that can achieve high information density, but more importantly, perform robust and reliable transcoding for a wide variety of data types in a cost-effective manner, is necessary for the development of DNA-based information storage in practical applications (24–26).

To achieve this goal, we propose the novel coding algorithm YYC, inspired from the traditional Chinese concept of “Yin-Yang”, which represents two different but complementary and interdependent rules, and we demonstrate its performance by simulation and experimental validation. The novelty of YYC is that the incorporation of “Yin” and “Yang” rules leads to a final 1,536 coding rules that can suit diverse data types and introduce encryption for data security. We have demonstrated that YYC can effectively eliminate the generation of long homopolymer sequences and keep the GC content of generated DNA sequences within acceptable levels. Two representative file formats were chosen to be stored as oligo pools *in vitro*, and a 54-kb DNA fragment was established in yeast cells for robustness evaluation. Our results show that YYC exhibits superior performance with regard to reliable data storage and physical information density reaching an EB/g scale.

## MATERIALS AND METHODS

### The Yin-Yang Codec strategy

#### Demonstration of the YYC transcoding principle

As an example (referred to as Rule No. 888 in Figure S1A), by the Yang rule, {A, T} represents the binary digit 0, and {G, C} represents the binary digit 1. By the Yin rule, the local nucleotide (current nucleotide to be encoded) is represented by the incorporation of the previous nucleotide (or “supporting nucleotide”) and the corresponding binary digit (Fig. S1B). During transcoding, the two rules are applied respectively for two independent binary segments and transcoded into one exclusive DNA sequence, and decoding occurs in reverse order. Taking the first nucleotide in Fig. S1B as example, the corresponding binary digits from segments “a” and “b” are 1 and 0, respectively. According to the Yang rule, the local nucleotide will have two options, {C, G}. In contrast, according to the Yin rule, the virtual supporting nucleotide is set as A by default, and thus the previous nucleotide A and the corresponding binary digit 0 will give the local nucleotide two options, {A, G} (Fig. S1A). The resulting encoded local nucleotide will be the intersection of the Yang rule, {C, G}, and the Yin rule, {A, G}, which leads to the consensus option {G}, and the process continues to the next binary digits. It should be noted that switching the binary segments will change the transcoded result, which means that {a: Yang, b: Yin} and {b: Yang, a: Yin} will result in the generation of completely different DNA sequences. A simple example of the transcoding demonstration process is illustrated in supplementary information (Fig. S1B, Supplementary video). Generally, only one fixed incorporated rule is selected for transcoding each dataset. Nevertheless, multiple rules can be used for transcoding for encryption purposes; the corresponding rule-usage information will be stored separately.

#### Incorporation of YYC transcoding pipeline

Considering the features of the incorporation algorithm, binary segments containing excessively imbalanced 0s or 1s tend to produce DNA sequences with extreme GC content or undesired repeats. Therefore, binary segments containing a high ratio of 0 or 1 (>80%) will be collected into a separate pool and then selected to incorporate with randomly selected binary segments with normal 0/1 ratios.

#### Constraint settings of YYC transcoding screening

In this study, a working scheme named YYC-screener is established to select valid DNA sequences. By default, generated DNA sequences (normally ~200-nt) with GC content greater than 60% or less than 40%, carrying > 6-mer homopolymer regions, or possessing a predicted secondary structure of < −30 kcal/mol are abandoned. Then, a new run of segment pairing will be performed to repeat the screening process until a generated DNA sequence meets all screening criteria. Considering that the sequencing and synthesis technologies continue to rapidly evolve, the constraint settings are designed as nonfixed features to allow user customization. In this work, the constraints are set as follows: GC content between 40% and 60%; maximum homopolymer length < 5; and free energy ≥ −30 kcal/mol (secondary structure free energy is calculated by Vienna RNA Version 2.4.6).

#### The theoretical information density estimation of YYC

In practice, denoting the original digital file as {0,1}*^n^*, the file will be transcoded by YYC into *x* oligos with the length *y* nucleotides (primer for amplification or tagging is not included). In addition, each oligo contains an error-correction code region at length of *z* bits. In addition, *δ* additional oligos for logical redundancy are set in YYC. Thus, the net information density *d* of YYC can be identified with the following formula:

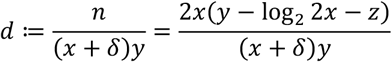

##### Lemma 1

the upper bound of its information density is:

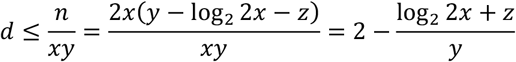

##### Lemma 2

when *δ* = *x*, the information density of YYC will reach its lower bound:

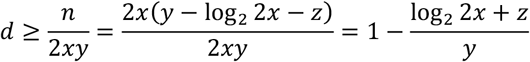

##### Proposition 1

Through the transcoding simulation, see Figure S3, if 100 rounds of random pair iteration are set in the encoding process, the open interval of 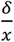 is (0,0.032). When 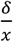 reaches 0.032, the lower bound is roughly equivalent to 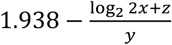. Suppose *x* = 30,000, *y* = 128, *z* = 8, with the range of 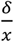 mentioned before, the range of information density is from 1.751 bits/nt to 1.814 bits/nt.

To further optimize the information density of YYC, two parameters can be adjusted based on digital data. As shown in Figure S4, replacing the rules in YYC can significantly reduce the production of additional oligos. Increasing random pair iterations can also achieve a similar effect.

Theoretically, in coding theory, storing binary data in the form of an oligo pool will introduce a few basic factors that will affect the information density. First, the oligo sequences are stored as a mixture of DNA molecules. Second, each oligo sequence is designed without long homopolymers or extreme GC content.

##### Definition 1

A code 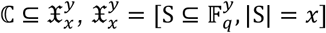, the largest code size is 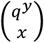. In particular, for DNA-based storage, 𝔽_q_is [A,T,G,C], and the information density 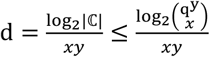.

##### Proposition 2

If *y* = 128, *x* = 30,000, and 𝔽_q_ = [A,T,G,C], then *d* ≤ 1.9897.

##### Lemma 3

A code 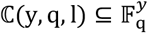, each codeword does not have a length of homopolymer runs greater than *l*, the largest code size *λ^n^* is the largest real root of the equation 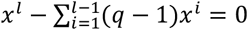, and the information density 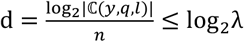.

##### Proposition 3

If *y* = 128, *1* = 4, and 𝔽_q_ = [A,T,G,C], then *d* ≤ 1.9757.

##### Definition 2

A code 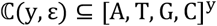, each codeword has [(0.5 – ε)y, (0.5 + ε)y] GC, the largest code size is 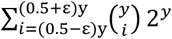, and the information density 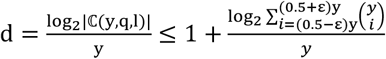.

##### Proposition 4

If y = 128 and *ε* = 0.1, then *d* ≤ 1.9997.

Based on the above, the information density *d* ≤ min[1.9897, 1.9757, 1.9997] = 1.9757, and the highest code information density d ≥ 2 - (2 - 1.9897) - (2 - 1.9757) - (2 - 1.9997) = 1.9651 (the corresponding settings include: generated DNA sequence length = 128, sequence number = 30,000, GC content between 40% and 60%, and homopolymer length ≤ 4).

### In-silico transcoding simulation

#### Computing and Software

All encoding, decoding, and error analyzing experiments were performed in a Windows 7 environment including an i7 GPU and 12 GB of RAM using Python 3.7.3 in the IDE PyCharm 2019.1.

#### Input files and parameters for simulation

The test files include 113 journal articles (including images and text), 112 mp3 audio files from Scientific American, and the supplementary video files from 33 journal articles.

To compare the compatibility of different coding schemes, all the test files were transcoded by Church’s code, Goldman’s code, Grass’ code, DNA Fountain, and YYC via the integrated transcoding platform “Chamaeleo” that we developed (27). The segment length of binary information was set as 32 bytes. For Church’s code, Goldman’s code, and Grass’ code, original settings as previously reported were used in this study. For DNA Fountain and YYC, the constraints were set as follows: GC content between 40% and 60%; max homopolymer allowed = 4. For free energy constraints in YYC, the threshold was set as ≥ −30 kJ/mol.

Additional transcoding simulation tests were performed to evaluate the robustness and compatibility of DNA Fountain and YYC. DNA Fountain source code was used to perform encoding and decoding tests on 9 different formats of files and 10 bitmap images with default parameter settings (c-dist = 0.1, delta = 0.05, header size = 4, redundancy = 7%, homopolymer = 4, GC: 40%-60%). The oligo length of both strategies was set as 152 bases with indices or seeds for data retrieval and without error-correction codes. To determine the minimum redundancy required for file decoding, a test interval of minimum redundancy was set as 1%, and the maximum redundancy allowed was 300%. In some cases, the process was terminated with a system error, which might be caused by stack overflow. The estimation result of iteration run, information density, and minimum required logical redundancy is summarized in Tables S3 and S4.

### Experimental validation by *in vitro* and *in vivo* data storage demonstration

#### File encoding using YYC and DNA Fountain

The binary forms of three selected files (9.26 ×10^5^ bits, 7.95×10^5^ bits, and 2.95×10^5^ bits) were extracted and segmented into three independent 128-bit segment pools. A 16-bit RS code was included to allow the correction of up to two substitution errors introduced during the experiment. Next, four 144-bit binary segments (data payload + RS code) were used to generate a fifth redundant binary segment to increase logical redundancy. Then, another 16-bit index was added into each binary segment to infer its address in the digital file and in the oligo mixture for decoding. Rule 888 from the YYC scheme was applied to convert the binary information into DNA bases. The aforementioned YYC-screener was used to select viable DNA sequences. Eventually, 8,087 160-nt DNA sequence segments were generated. To allow the random-access of each file, a pair of well-designed 20-nt flanking sequences was added at both ends of each DNA sequence. Finally, an oligo pool containing 10,103 single-stranded 200-nt DNA sequences was obtained.

For DNA Fountain, the recommended default settings from its original report (23)(c-dist = 0.1, delta = 0.5, header size = 4, homopolymer = 4, GC: 40%-60%), with the exception of redundancy, were used to generate DNA oligo libraries. To ensure successful decoding, minimum redundancy was determined. Therefore, 13%, 22%, 73%, and 12% logical redundancy was added for a tar-archiving compressed file, text1, text2, and image files, respectively. Finally, an oligo library encoding a tar-archiving compressed file (9,185 sequences) and an oligo library encoding the mixed three individual files (10,976 sequences) were obtained.

A part of one text file, ~13 kB, was transcoded into DNA sequences by YYC for *in vivo* storage using a similar procedure, but the binary segment length was set as 87 bytes (or 456 bits). As described in the main text, the sequence was divided into 113 data blocks with 456-nt each. To increase the fidelity, a 24-nt RS code was added. The total data payload region as double-stranded DNA for *in vivo* storage is (456+24) × 113 = 54,240 bp.

#### Synthesis and assembly

The three oligo pools were outsourced for synthesis by Twist Biosciences and delivered in the form of DNA powder for sequencing.

For *in vivo* storage, the 54,240-bp DNA fragment was first segmented into twenty sub-fragments (2,500 – 2,900 bp) with overlapping regions and then further segmented into building blocks (800 – 1,000 bp, hereafter referred to as blocks). For each block, twenty 80-nt oligos were synthesized with a commercial DNA synthesizer (Dr. Oligo, Biolytic Lab Performance, Inc.) and then assembled into blocks by applying the polymerase cycling assembly (PCA) method using Q5 High-Fidelity DNA Polymerase (M0491L/NEB) and cloned into an accepting vector for Sanger sequencing. Then the sequencing-verified blocks were released from their corresponding accepting vector by enzymatic digestion for the assembly of sub-fragments by overlap extension PCR (OE-PCR). Gel purification (QIAquick Gel Extraction Kit/28706/QIAGEN) was performed to obtain the assembled sub-fragments. By transforming all 113 sub-fragments (300 ng each) and the low-copy accepting vector pRS416 into BY4741 yeast using LiOAc transformation (28) and taking advantage of yeast’s native homologous recombination, the full-length ~54-kb DNA fragment was obtained. After 2 days of incubation on selective media (SC-URA/630314/CLONTECH) at 30°C, three single colonies were isolated for liquid culturing in YPD (Y1500/SIGMA) before sequencing.

#### Library preparation and sequencing

For library preparation of the synthesized oligo pool, the DNA powder was first dissolved in ddH_2_O to obtain a standard solution (SS) with an average 10^6^ molecules/μl for each synthesized oligo pool. Then the SS was diluted 100, 1,000, and 10,000-fold to create the three working solutions (WSs) WS0, WS1, and WS2 with average concentrations of 10^4^, 10^3^, and 10^2^ DNA molecules/μl, respectively, for each oligo pool. Then each WS was amplified by PCR with three technical replicates to obtain the amplified product for P2 and each of the three different files for P1 and P3. PCR amplification was performed using KAPA HiFi HotStart Ready Mix PCR kit (Roche KK2602/07958935001); 2 μL forward and reverse primer pairs (10 μM each), 1 μL template DNA, and 30.5 μL ddH_2_O were added to a final reaction volume of 50 μL. The PCR thermal cycler program settings were as follows: 98°C for 5 min; 20/23/26 cycles of 98°C for 10 s, 55°C for 15 s, and 72°C for 10 s; and a final extension of 72°C for 2 min. All amplified DNA libraries were then sequenced using DIPSEQ-T1 sequencing (29).

For *in vivo* storage, the methods for genomic DNA extraction and standard library preparation of the 12 selected yeast colonies were described in previous studies. The prepared samples were sequenced using the DNBSEQ-G400 (MGISEQ-2000) sequencing platform (30).

#### Data analysis

In total, 2.76 G PE-100 reads were generated for the *in vitro* storage experimental validation. Sequencing data with an average depth of 100x were randomly subsampled for information retrieval. The reads were first clustered and assembled to complete sequences for each type of oligo. Flanking primer regions were removed, DNA sequences were decoded to binary segments using the reverse operation of encoding, and substitution errors were corrected using RS code. The binary segments were re-ordered according to the address region. During this process, additional information was removed based on the address. The complete binary information was then converted to a digital file. The data recovery rate was calculated using 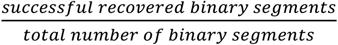 (Tables S5 and S6). For error analysis, sequencing data with average depths of 100x, 300x, 500x, 700x, and 900x were randomly subsampled 6 times using different random seeds.

The probability of successful amplification *P* can be calculated by the following formula with the hypothesis that each type of DNA sequence has identical copy number:

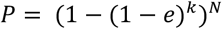

where *e* is the amplification efficiency, *k* is the copy number per molecule, and *N* represents the number of types of DNA sequences. When *N* increases, *P* will decrease accordingly. When *k* increases, *P* will increase accordingly. The amplification efficiency for ~50-kb circular DNA is ~85% (31), whereas the PCR amplification efficiency with an ~0.5-kb linear template is ~95%. Therefore, the probability of successful amplification of the 54-kb circular DNA used in our work is 97.75%. With identical molecule copy number, if storing the information as 113 480-nt fragments, the probability of successful amplification is 75.36%. If the total molecules of 113 fragments with average copy number of 4 and 5 follow a Poisson distribution (32), then the probability of getting all the DNA fragments with more than one copy are only ~12% and ~48% respectively (calculation based on 10 independent repeat simulation results of 10,000 iterations of random sampling). In this case, even if the amplification efficiency is 100%, the maximum odds to get all the target sequence to be amplified cannot reach 100%.

In total, 47.6 M PE-100 reads were generated for *in vivo* storage, in which the 10% low-quality reads (Phred score < 20) by SOAPnuke were filtered (33). Reads of the host genome were removed using samtools after mapping by BWA (34–36). Short reads were then assembled into contigs by SOAPdenovo (37,38). Blastn was used to find the connections between contigs (39). A python script was written to merge the contigs and obtain the assembled sequences for each strain. Multiple sequence alignment (MSA) was conducted to align the assembled sequences by clustalW2 for majority voting process to identify structural variations, insertions, and deletions (40). Pre-added RS codes were used for error correction of substitutions. The complete DNA sequence was decoded by the reversing the operations of encoding to recover the binary information.

The *in vivo* information density is calculated by:

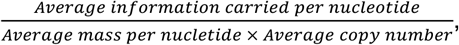

where:

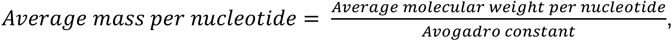

and:

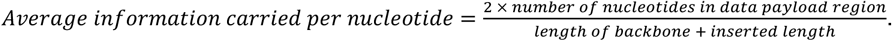

The average molecular weight per nucleotide is 330.95 g/mol, the average copy number is 2, the length of data payload region is 51,528 bp, the backbone size of pRS416 is 4,898 bp, and the inserted length is 54,240 bp. Therefore, the information density is calculated to be 1.59 × 10^21^ bits per gram of DNA, which equals 1.988 × 10^20^ bytes per gram of DNA.

## RESULTS

### General principle and features of the Yin-Yang codec

In nature, DNA usually exists in a double-stranded structure. In some organisms such as phages, both strands encode genetic information to make the genome more compact. Inspired by this natural phenomenon, we used the basic theory of combinatorics and cryptography and developed a new codec algorithm on the basis of Goldman’s rotating encoding strategy (41,42). The codec system we developed here can successfully address the challenges of generating DNA sequences with long homopolymers, extreme GC content, or complex secondary structures, which introduce difficulties for DNA synthesis and sequencing. The codec provides dynamic combinatory rules and thus can generate optimal DNA sequences for a particular file compared to early coding schemes using fixed mapping rules.

The general principle of the YYC algorithm is to incorporate two independent encoding rules, “Yin” and “Yang”, into one DNA sequence (referred to as incorporation hereafter), thereby “compressing” two bits into one nucleotide. As shown in Fig. 1A, N1/N2/N3/N4 represent the different nucleic acids A/T/C/G. When n is an integer chosen from 1 to 8, X_n_+Y_n_=1 and X_n_×Y_n_=0. Under one selected combinatory rule, an output DNA sequence is generated by the incorporation of two binary segments of identical length. In the Yang rule, this can provide 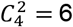 different rules. In the Yin rule, N1 and N2 are mapped to different binary digits, while N3 and N4 are also mapped to different binary digits independent of N1 and N2. This ensures that for one position, the Yin and Yang rules will have one and only one consensus base (Fig. S1 and Supplementary Video). Meanwhile, according to the four different options for the previous nucleotide, the two groups (N1/N2 and N3/N4) also have independent options for mapping to 0 and 1. Therefore, the incorporated “Yin” and “Yang” rules will provide a total of 1,536 (6×256) combinations of transcoding rules to encode the binary sequence (Fig. 1A).

**Figure 1.**
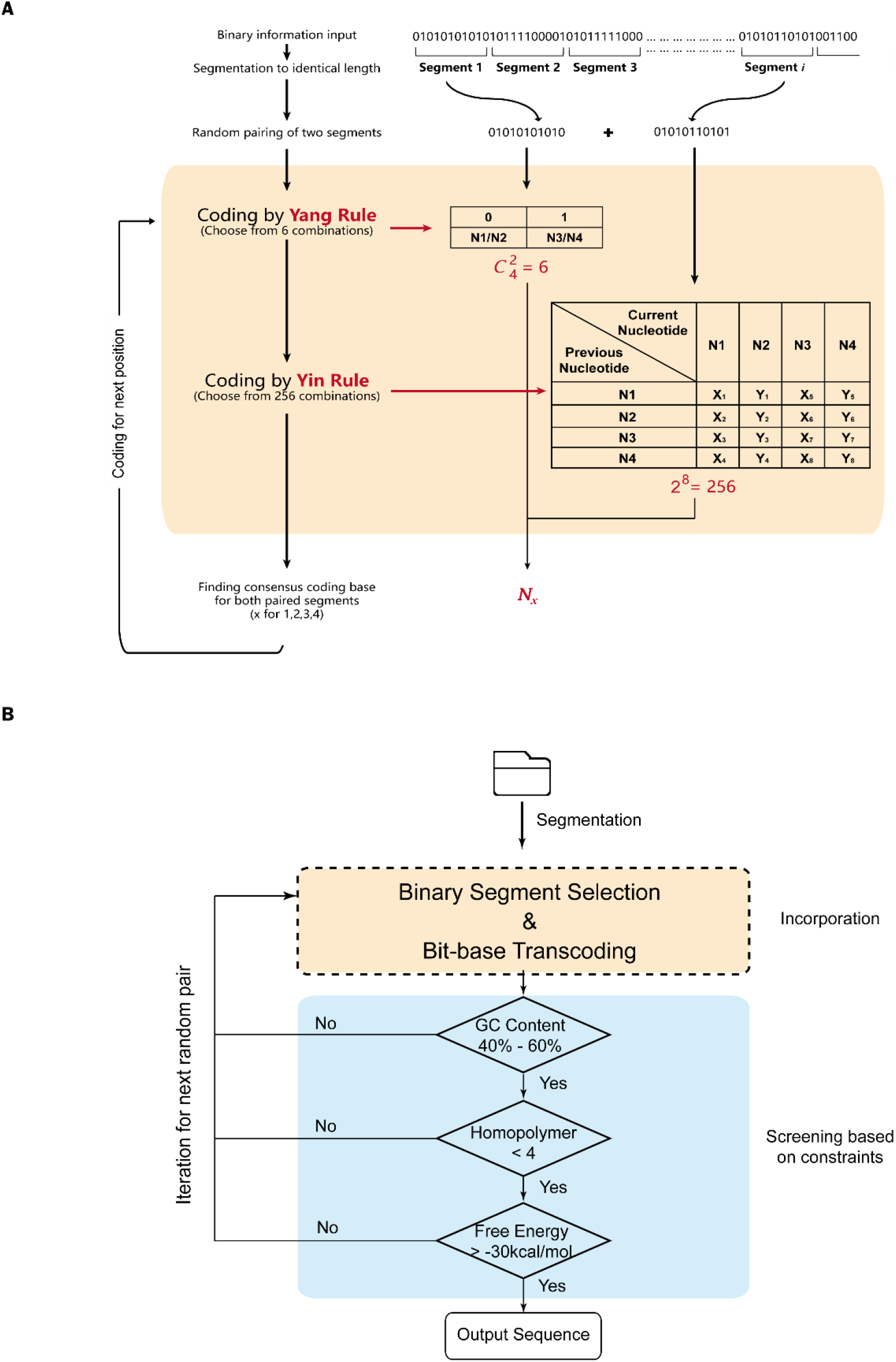
Principles of the Yin-Ying Codec. A) The bit-to-base transcoding process of YYC. N1/N2/N3/N4 represent different nucleic acids A/T/C/G. When n is an integer chosen from 1 to 8, X_n_+Y_n_=1, and X_n_×Y_n_=0. B) The flowchart of the YYC encoding pipeline.

The pipeline for the YYC system includes three main steps: segmentation, incorporation, and validity-screening (Fig. 1B). For segmentation, the binary information is extracted from a source file and partitioned into multiple segments according to requirements. Indices are added to each segment, and a pool of binary segments of identical length are then obtained. In the incorporation step, two segments are selected randomly to be incorporated into one DNA sequence according to the YYC algorithm. For the validity-screening in the last step, the incorporated sequence will be subjected to screening against pre-set constraints, including GC content, maximal homopolymer length, the secondary structure free energy, etc., and only sequences that meet set criteria will be considered as valid sequences. If the sequence fails to meet these criteria, the iteration process will be activated. Another randomly generated binary segment from the library will be added for incorporation with the first selected segment until a valid sequence is generated. Occasionally, when most of the binary segment contains more than 80% repeated 0s or 1s, the encoding process may enter an infinite loop or generate a poor result. Therefore, we established a “firewall” that sets the upper limit of iterations to 100 to avoid this situation (Fig. S2, Table. S2).

As estimated previously, the greatest information density for DNA storage is 2 bits per base. However, this density cannot be realistically attained due to intrinsic biochemical constraints and technical limitations of the DNA synthesis/sequencing procedures (43). In addition, indices for providing the corresponding address in the library and the introduction of an error correction code for decoding robustness and PCR amplification primers for random access of information in the DNA sequence will further reduce the information density. For previously established coding algorithms, the information density is a fixed value. It is either close to the theoretical value with associated risk of transcoding failure, or it is a relatively low level and thus compromises cost efficiency. In contrast, for YYC, the information density of each transcoding rule among the 1,536 options is different, and a maximum 1.965 bits per base can be attained under specific constraints (see Materials and Methods & Fig. S3). Such a design scheme allows the user to choose an optimal trade-off between robustness and cost efficiency in dealing with files showing specific binary data patterns (Table S1). This design also holds the opportunity that when current technical bottlenecks in synthesis and sequencing procedures are resolved, extreme information density can be achieved.

To demonstrate the compatibility of the YYC algorithm and quantify its featured parameters in comparison to other early-established DNA-based data storage coding schemes, a collection of different formats of files including text, image, audio, and video files, with a total size of ~1 GB, was transcoded by YYC and other early established coding algorithms for comparison (1,21–23). A focus was placed on the features of generated DNA sequences, including the GC content, the maximum homopolymer length, and the free energy of the secondary structure. For most coding schemes, the GC content of generated DNA sequences shows a relatively wide range from 0% to 100%. In contrast, the GC content of sequences generated by YYC and DNA fountain is largely between 40% to 60% (Fig. 2A). The reason for this difference is that the incorporation of binary segments for both YYC and DNA fountain is flexible, which provides more possibilities for obtaining DNA sequences with desired GC content. Like all other coding algorithms, YYC also introduced the screening constraints to avoid the generation of long homopolymers, and the maximum homopolymer length is set at 4 in consideration of computing resources as well as the technical limitations of DNA synthesis and sequencing. In addition, YYC also takes the secondary structure of generated DNA sequences into consideration as part of compatibility by rejecting all DNA sequences with free energy lower than −30 kcal/mol (Fig. 2B).

**Figure 2.**
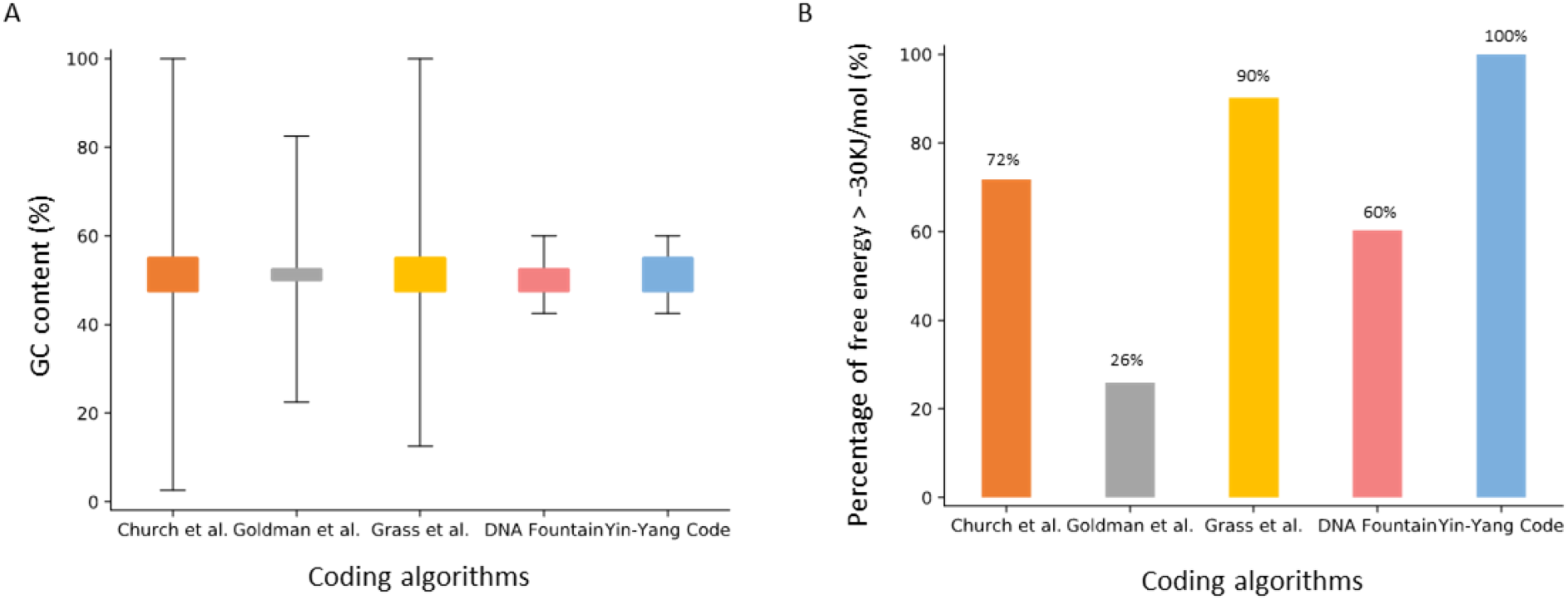
The compatibility of generated DNA sequences using different coding schemes. A) Distribution of GC content of generated DNA sequences using different coding strategies. B) Percentage of generated DNA sequences with free energy for predicted secondary structure > −30 kJ/mol using different coding strategies. The length of the DNA sequence is set at 128 nt for the analysis.

Given the above, YYC offers the opportunity to generate DNA sequences that are highly amenable to the “writing” (synthesis) and “reading” (sequencing) processes. This is crucially important for improving the practicality and robustness of DNA data storage.

### The robustness of YYC for stored data recovery

For DNA data storage, robustness is primarily affected by errors introduced during “writing” and “reading”. There are two main types of errors: random error and systematic error. Random errors are often introduced by the synthesis or sequencing error of a few DNA molecules, and this can be redressed by mutual correction with increased sequencing depth. Systematic errors refer to observed mutations in all DNA molecules, including insertions, deletions, and substitutions, which are introduced during synthesis and PCR amplification (referred to as common errors), or the loss of partial DNA molecules. In general, it is difficult to correct systematic errors, and they will thus lead to the loss of stored binary information to varying degrees.

Compared with substitutions, insertions and deletions will change the length of the data-encoded DNA sequence and introduce challenges to the decoding process. To test the robustness of YYC against common errors, we randomly introduced the three most commonly seen errors into the DNA sequences at a rate of 0.3% for each type and analyzed the corresponding binary information recovery rate. Compared with other coding algorithms excepting DNA fountain, YYC exhibits equivalent robustness for data recovery in the presence of substitutions without the use of error correction; the data recovery rate is > 99% (Fig. 3A). A similar trend is also observed in the presence of insertions and deletions without the use of error correction (Fig. 3C). This difference between DNA fountain and other algorithms including YYC occurs because uncorrectable errors can affect the retrieval of other data packets through a domino effect for DNA fountain. The introduction of error-correction codes is universally acknowledged as an effective strategy to correct common errors at a certain level (22,23,44). We also evaluated each coding algorithm’s binary information recovery rate in addition to the introduction of error-correction codes. Our results suggest that an error-correction code helps to fix the substitution and improve the average recovery rate by 72.57% for DNA Fountain and 0.31% for other coding schemes including YYC (p < 0.01, Fig. 3B). In contrast, traditional Reed-Solomon (RS) error-correction codes poorly fix insertion and deletion errors and thus cannot improve the recovery rate (p > 0.05, Fig. 3D). Although researchers have been aware of this issue and have developed a few specific error-correction codes, these strategies need more experimental verification (45–47). Currently, sequences with insertions or deletions will be treated as lost sequences.

**Figure 3.**
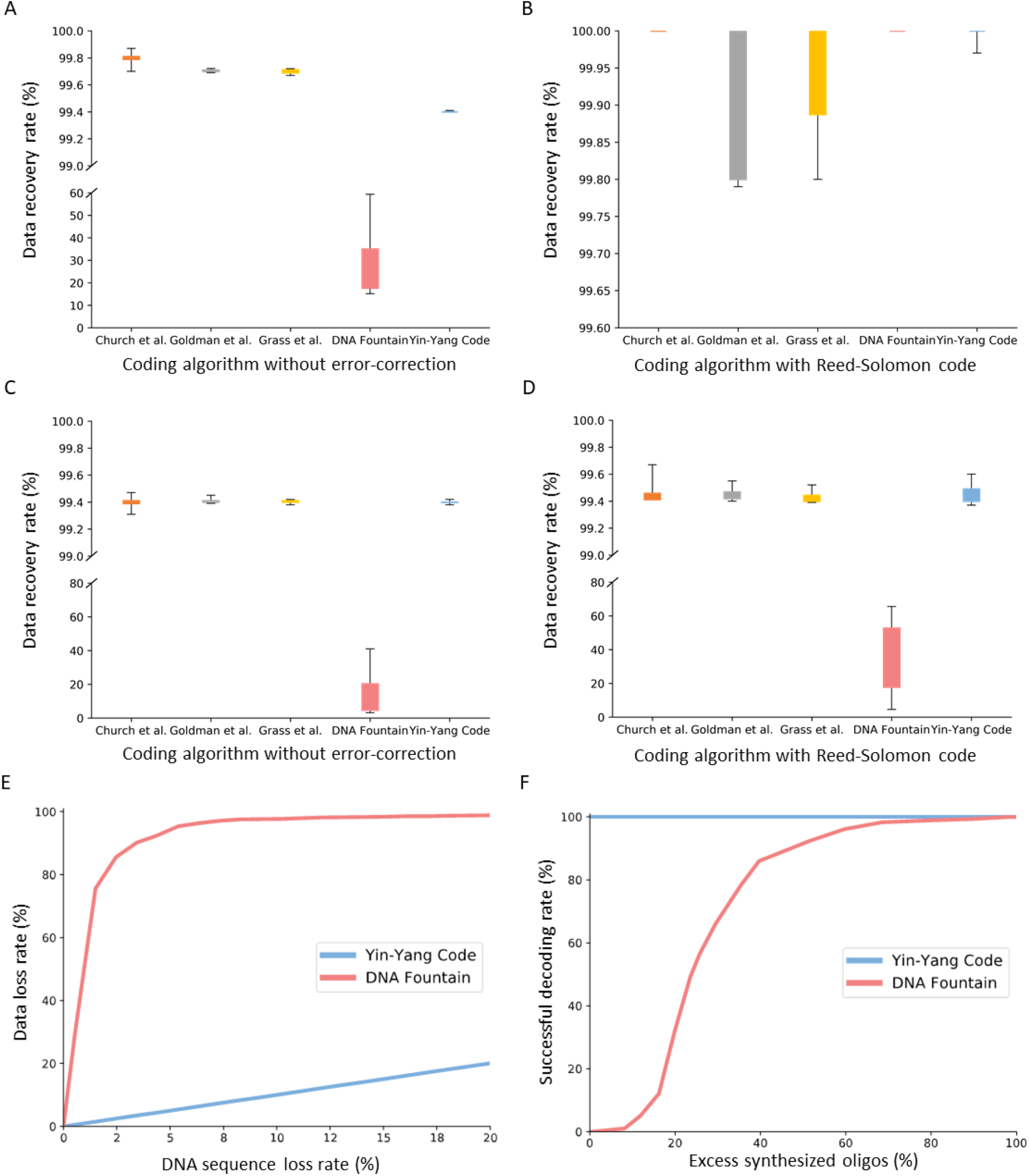
Robustness analysis for different coding schemes. The binary data recovery rate of different coding strategies without error-correction algorithm for A) randomly introduced substitutions and C) insertions/deletions in comparison with the data recovery rate with error-correction algorithms for B) randomly introduced substitutions and D) insertions/deletions. For B) and D), 2 bytes of RS code were introduced. The Y axis represents the percentage of successful recovered binary segments from the total number of stored binary segments. E) The binary data loss rate evaluation of Yin-Yang and DNA Fountain coding strategies under a simulated DNA sequence loss rate gradient. F) The success rate of decoding with Yin-Yang Codec and DNA Fountain under a simulated gradient of excess synthesized oligos.

It is also important to establish mechanisms to avoid total data retrieval failure caused by the loss of partial DNA molecules (48). Like early coding schemes (Church et al., Goldman et al., Grass et al., etc.), YYC is also designed as linear block non-erasure codes, which show a linear relationship between data loss and encoded sequence loss. Nevertheless, because of the convolutional binary incorporation of YYC, errors that cannot be corrected within one DNA sequence will lead to the loss of information for two binary sequences. In contrast, DNA fountain uses a different data retrieval strategy based on its grid-like topology of data segments, and theoretically its data recovery cannot be guaranteed when a certain number of DNA sequences are missing (23). In our study, the *in-silico* calculation of the information recovery rate in the context of a gradient of DNA sequence loss was performed. Our results show that YYC is capable of maintaining linear retrieval as predicted. The data recovery percentage can be maintained at > 97% when the sequence loss rate is < 2%. Even with 20% sequence loss, YYC can recover the remaining ~80% of data (Fig. 3E). In contrast, the DNA fountain coding scheme exhibits better performance compared to YYC when sequence loss is < 1.8%. However, when the sequence loss rate exceeds 2%, the data recovery rate becomes highly volatile and drops significantly. Only ~2% of the information can be recovered when the sequence loss rate exceeds 10% (Fig. S4).

In addition, information loss and decoding failure in DNA data storage can also originate from original defects in the transcoding algorithms. Theoretically, erasure codes such as the Luby Transform (LT) code risk the possibility of losing original information during encoding because the encoded matrix does not necessarily satisfy the full rank condition for decoding information with particular binary features (49–51). Meanwhile, an erasure channel may lead to further loss of original information and increase the probability of decoding failure (25,52). According to a previous study by Lu et al., a formula was developed to analyze the probability of decoding failure *f* when 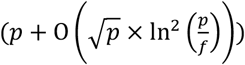 data packets are received, in which *p* represents the number of original packets (25). Because the formula refers to logical redundant packets added in addition to original packets *p*, if *f* needs to be maintained at a low level, the logical redundancy must increase (52). Therefore, increasing logical redundancy by synthesizing excessive DNA oligos encoding extra information could greatly improve the probability of successful decoding for DNA Fountain. However, this has no effect on YYC (Fig. 3F). Further transcoding simulation has demonstrated that for uncompressed file formats, DNA fountain demonstrated variable requirements for the level of logical redundancy, which leads to a varying information density (Tables S3 & S4). Considering that the operations of DNA are not as convenient as those of digital data and that DNA synthesis and data retrieval are “heterochronic” in real applications, which makes the immediate additional transmission of data packets impossible, coding schemes risking uncertain decodability are not ideal for DNA-based data storage applications.

### Experimental validation of the compatibility of YYC

To determine whether the real performance of YYC shows agreement with the previous simulation, we performed experimental validation by encoding three digital files including two text files, one each in English and Chinese, and an image using YYC and storing the transcoded file in the form of 10,103 200-nt oligos (*in vitro*) and an ~54-kb DNA fragment composed of 113 480-bp data blocks in (*in vivo*) (53) (Fig. 4A). The sequence design of YYC transcoding-generated oligos includes 20-nt primers at the 5’ and 3’ ends, a 128-nt data payload region, a 16-nt RS code region, and a 16-nt index region. The sequence design of the 480-bp data block includes the 456-bp data payload region and the 24-nt RS code region (Fig. 4B).

**Figure 4.**
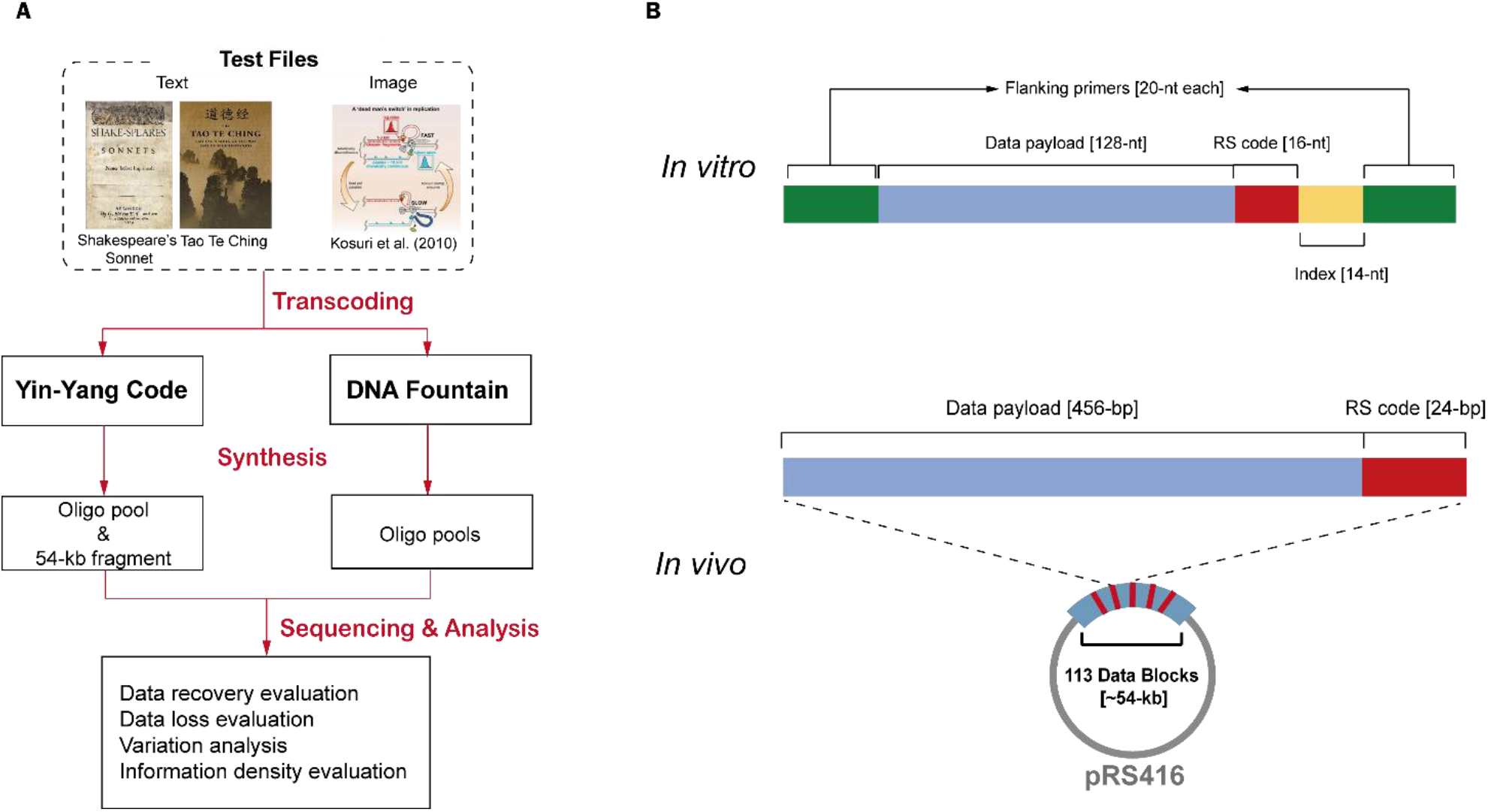
The validation experiment design and analysis workflow for *in vitro* and *in vivo* binary data storage using Yin-Yang Codec and DNA Fountain coding strategies. A) The experimental validation workflow including binary information transcoding, synthesis of generated DNA sequences, and robustness analysis using generated sequencing to access stored binary data. B) The design of DNA sequences generated by Yin-Yang Codec for *in vitro* data storage in the form of 200-nt oligonucleotides and *in vivo* data storage in the form of a 54-kb DNA fragment inserted in the low-copy plasmid pRS416.

For *in vitro* storage validation, 25% logical redundancy (i.e., 4 sequences will generate 1 logically redundant sequence) was introduced into the 10,103 oligos (P1) in case of sequence loss. Meanwhile, to compare the robustness, two independent oligo pools of these three files transcoded by DNA fountain were also synthesized (23): One oligo pool (P2) used a tar-archiving compression package with 13% logical redundancy and includes 9,185 oligos, and the other oligo pool (P3) encoded these three files individually with minimum logical redundancy suggested for successful decoding and includes 10,976 oligos. The sequence structure of oligos is designed according to the principles described in the original study.

The average molecule copy (AMC) number of the commercially synthesized P1, P2, and P3 pools is estimated at 1.58×10^8^, 1.92×10^8^, and 1.54×10^8^, respectively. A 10-fold serial dilution of P1, P2, and P3 was performed for sequencing to test the minimal copy number of oligos required for successful file retrieval and robustness against DNA molecule loss. Working solutions with estimated AMC numbers of 10^4^, 10^3^, and 10^2^ for each oligo pool were obtained after serial dilution. The sequencing results demonstrate that > 99.9% of the corresponding data from P1 can be recovered at AMC numbers of 10^4^ and 10^3^, with no preference for a specific data format (Fig. S5). The recovery rate shows an increase of instability, varying from 55.45% to 87.53% at the AMC number of 10^2^. For DNA fountain, the recovery rate for the AMC number of 10^4^ is comparable to that of YYC and drops significantly at AMC numbers of 10^3^ and 10^2^ with average recovery rates at 22.87% and 0.33%, respectively (Fig. 5A). In particular, for P2, the data was first tar-archived and then transcoded for storage, and a maximum 32.83% of the data package can be retrieved. However, the disruption of the compression package leads to the total loss of original data. The sequencing result also demonstrates that the relationship between the information loss rate and the sequence loss rate of each synthesized oligo pool exhibits consistency with the rates in in-silico simulations for YYC and DNA fountain (Fig. 5B). It has been suggested previously that most random errors introduced during synthesis or sequencing can be corrected by increased sequencing depth (54). Our result shows that although lost sequences could be retrieved by deep sequencing (Fig. S6A), these sequences are at relatively low depth, and they contain more errors compared with other sequences (Fig. S6B). Therefore, the retrieval of these types of sequences makes them insufficient for valid information recovery. Our result suggests that the loss of DNA molecules is the major factor affecting the data recovery rate, and even high sequencing depth could not improve the recovery rate if a certain quantity of data-coding DNA molecules is lost.

**Figure 5.**
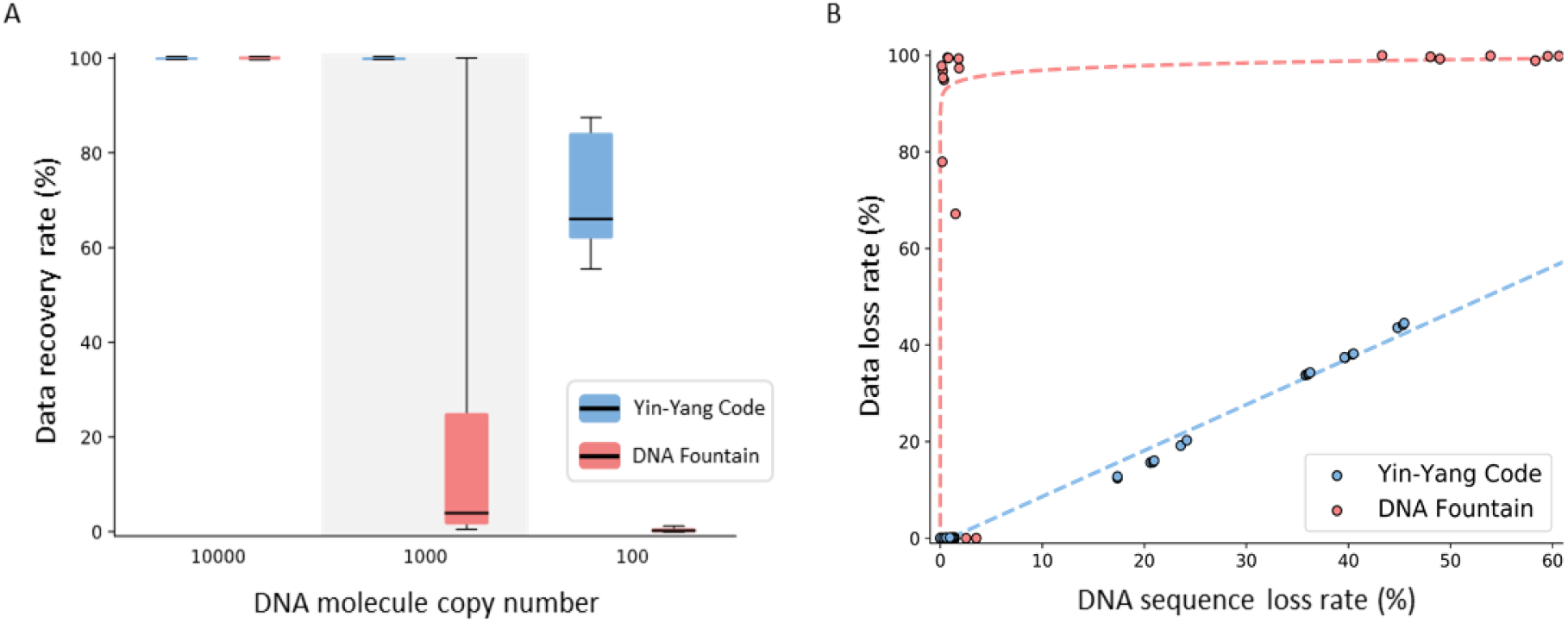
Robustness analysis of *in vitro* binary data storage by Yin-Yang Codec and DNA fountain. A) The binary data recovery rate of both strategies at different copy number of DNA molecules. B) The binary data loss rates of both strategies at different DNA sequence loss rates.

*In vivo* DNA data storage has drawn attention in recent years for its potential to enable economical write-once encoding with stable replication for multiple data retrievals, and the data stability during transmission over generations has also been validated (44). Meanwhile, the random access of data does not rely on the specific designed amplification primers, and thus the data payload region can be further increased to allow higher information density. In addition, according to our estimation, storing the binary information as one large fragment has the advantage of a higher possibility of data recovery compared to DNA molecule mixtures containing several smaller fragments. The *in vitro* amplification efficiency of a 50-kb DNA fragment using previously reported technique is ~85% (31), whereas that of a 500-mer DNA fragment is ~95%. If the copy number of each form of DNA molecule is 2, then the probability of complete successful amplification for the 50-kb DNA molecule (~99.75%) will be much higher compared to a hundred 500-mer DNA molecules (~75.36%), which contain an equal quantity of binary information. In real experiments, the synthesized DNA molecules as a mixture follow a Poisson distribution (32), which makes the probability of successful amplification of all DNA molecules even lower. Therefore, we transcoded a portion of one of the text files (Shakespeare Sonnet.txt) into a 54,240-bp DNA fragment using YYC and evaluated its potential in data robustness and information density for the application of *in vivo* DNA data storage. The generated DNA fragment was *de novo* synthesized and assembled into the low-copy centromeric plasmid pRS416 in yeast strain BY4741. To retrieve the stored information, we randomly selected 3 single colonies from the resulting yeast cells for whole genome sequencing and *de novo* assembly of sequencing reads to generate the full DNA sequence. Compared with the designed sequence, we identified in total 66 single-base substitutions in the construct. All of the designed sequence was first synthesized and assembled into 3-kb sub-fragments and verified by sequencing before full-length assembly in yeast. Our result suggests that the substitutions represent residual homologous recombination “patchworks” and could be corrected by the introduced RS code in the data blocks (Table S7). By doing so, we have demonstrated that the stored file can be perfectly retrieved. In addition, because the copy number of pRS416 at approximately 1 to 3 copies also represents a physical redundancy of stored information, we estimated the corresponding information density at ~198.8 EBytes/gram on average, which is three orders of magnitude higher than the results in prior work (23,55,56).

## DISCUSSION

The superiority of the YYC transcoding algorithm includes several aspects. First, it successfully balances high robustness, compatibility, and considerable information density for DNA data storage compared with other early efforts. With the gradual popularization of DNA data storage, it is crucially important that the developed coding algorithm can perform robust and reliable transcoding for a wide variety of data types, especially for data with specific binary patterns. Although specific binary patterns may be excluded by using compression algorithms, such as Lempel–Ziv–Welch (LZW), Gzip and run-length encoding (RLE), prior to transcoding to improve the compatibility of generated DNA sequences. However, because compression will change the original information structure, our result shows that even a partial loss of DNA molecules will end with the total failure of compressed data recovery. Current compression algorithms are not designed for DNA data storage, and further refinement can be performed to compress data appropriately for robust bit-to-base transcoding. Another potential advantage for YYC is the flexibility of rule incorporation from its 1,536 options. In our current workflow, the random incorporation of rules is used during the screening process to obtain valid DNA sequences according to constraints. In consideration of broader application scenarios, for example the need to store classified files with protection mechanisms for data confidentiality, YYC offers the opportunity to incorporate multiple rules for one file transcoding and thus provide a novel strategy for secure data archiving.

For the *in vitro* DNA storage demonstration in our study, the cost per base of oligo pool synthesis is approximately 1.2×10^-3^ US dollars, which equals an investment of $4.8 million per Gb of data to be stored in DNA. The input is even higher for DNA fountain because it requires a certain level of excessive DNA oligos encoding extra information to ensure high probability of successful decoding, at the cost of decreasing the information density and increasing the cost of DNA synthesis. Our sequencing results show that the error rate of synthesized oligo pools is ~ 1%, and in addition, ~1.2% of oligos are lost when mapping with the designed oligo sequence collections. There are two main steps in which errors could be introduced in our study: the synthesis of data-encoded oligos, and PCR amplification to obtain a sufficient amount of DNA for sequencing. In general, random errors introduced by sequencing can be easily corrected by sufficient sequencing depth, but errors introduced by PCR amplification can be problematic. Error-correction codes can improve information retrieval, but logically redundant sequences can play a more important role in retrieving lost sequence and correcting errors compared with error-correction codes for reliable DNA-based data storage. Another optional strategy is using both inner and outer error correction codes to recover some sequence loss as reported in previous work (22). In the future, DNA synthesis technology with high throughput and fidelity, which could yield a high quantity of DNA and avoid amplification, would be able to address this issue. For the *in vivo* DNA storage demonstration, we also find that there are random 1-nt indels and deletions with varying sizes from 5 kb to over 10 kb in different data coding regions across the selected three single colonies. These types of variations can be filtered out during the *de novo* assembly of sequencing reads, and they do not affect the data recovery process. However, the random variation occurring during the generation of host cells might also affect the data robustness in the form of *in vivo* storage, and thus future efforts to improve the stability of exogenous artificial DNA in host cells is necessary. Compared to another recent study of data storage using artificial yeast chromosomes in living yeast cells, our result shows better performance in information density and cost-effectiveness (44).

To summarize, we reported the establishment of a robust transcoding system named YYC, which can generate DNA sequences that are more compatible with current writing and reading technology without greatly sacrificing information density. We performed *in silico* simulations with a collection of different types of files including text, image, audio, and video. Our results suggest that YYC possesses outstanding compatibility and robustness compared with previously reported algorithms. Furthermore, experimental validation by both *in vitro* and *in vivo* storage further demonstrated YYC’s potential to be used for DNA data storage with broader applications.

## Supporting information

suppementary video 1

## DATA AVAILABILITY

The code package for YYC is available in the GitHub repository (https://github.com/ntpz870817/DNA-storage-YYC).

The data that support the findings of this study have been deposited in the CNSA (https://db.cngb.org/cnsa/) of CNGBdb with accession code CNP0001650.

## SUPPLEMENTARY DATA

Supplementary Data are available at NAR online.

## ACKNOWLEDGEMENTS

We thank Professor Craig Hunter and Dr. Chun-Ting Wu from Harvard University, and Professor Gennian Ge from Capital Normal University for constructive discussions and useful suggestions. We are thankful for support on DNA fragment synthesis and assembly for *in vivo* storage from China National GeneBank (CNGB).

## AUTHOR CONTRIBUTIONS

Y. S., Z.P., S.C., G. Z., and X.H. designed the experiment. Z.P. and S.C. conducted simulation and data analysis. G.Z. conducted the sequencing data analysis. S.J.Z. and H.Z. wrote and improved the code of the software program. J.C. and T.C. conducted the in vivo DNA fragment assembly. Z.L., H.Z., and Z. P. conducted the theoretical justification. Z.P. and Y.S. drafted the manuscript. Z.P., H.Z., and Y.S. prepared the figures and tables. G.Z., S.J.Z., H.H.L, G.C., and Y.S. revised the manuscript. H.Y., X.X., G.C., and Y.S. supervised the study. All authors read and approved the final manuscript.

