## supplementary file for "Towards Practical and Robust DNA-Based Data Archiving Using ‘Yin-Yang Codec’ System"

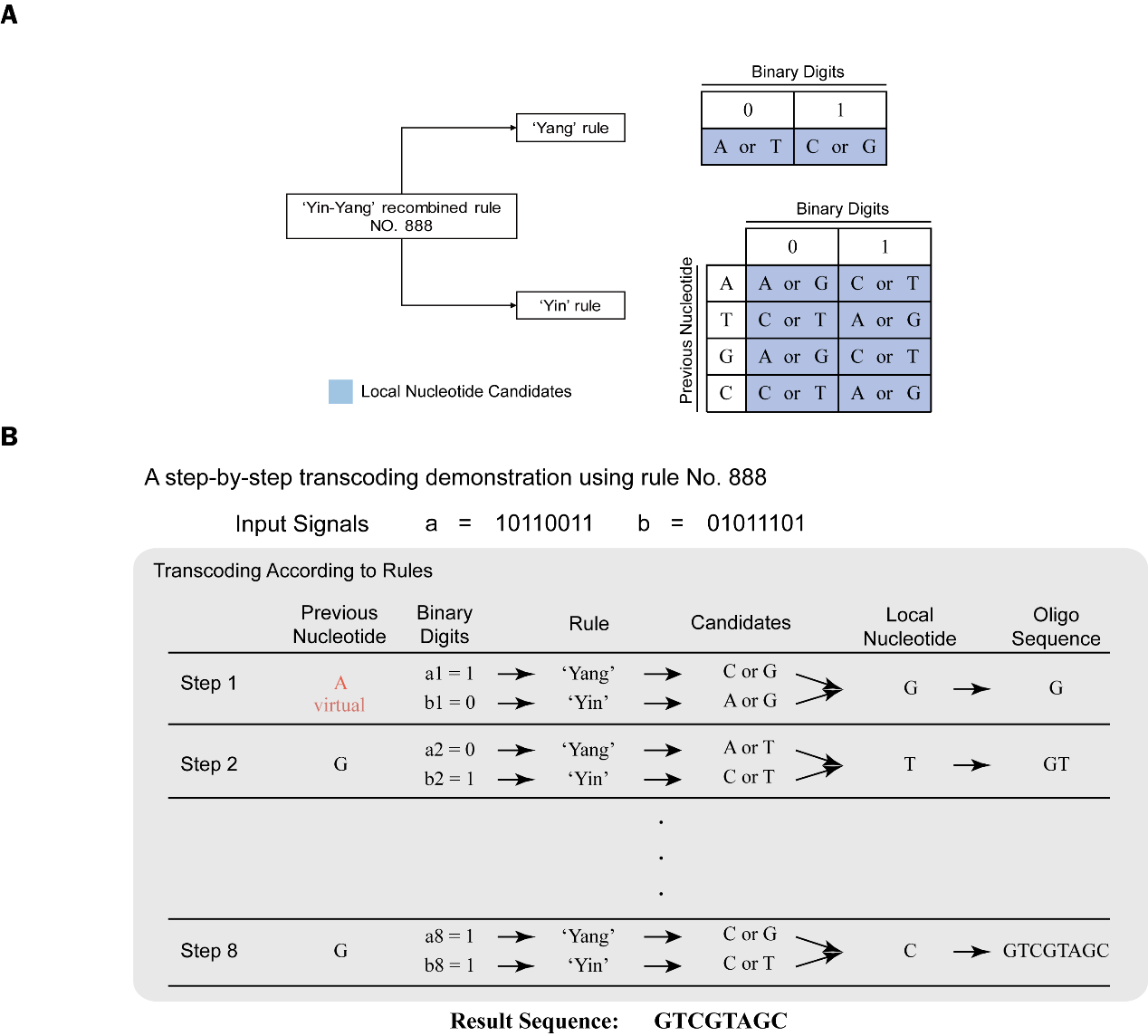

**Figure S1. The transcoding illustration using one of the 1,536 rules of YYC.** (A) The coding principle of YYC rule No. 888. (B) The demonstration of step-by step YYC transcoding process. ‘a1’ and ‘b1’ represent the first-position binary digit in segments ‘a’ and ‘b’, and so on. Virtual base A means this base is used only for determining the output base in the first run of transcoding and will not appear in the resulting sequence.

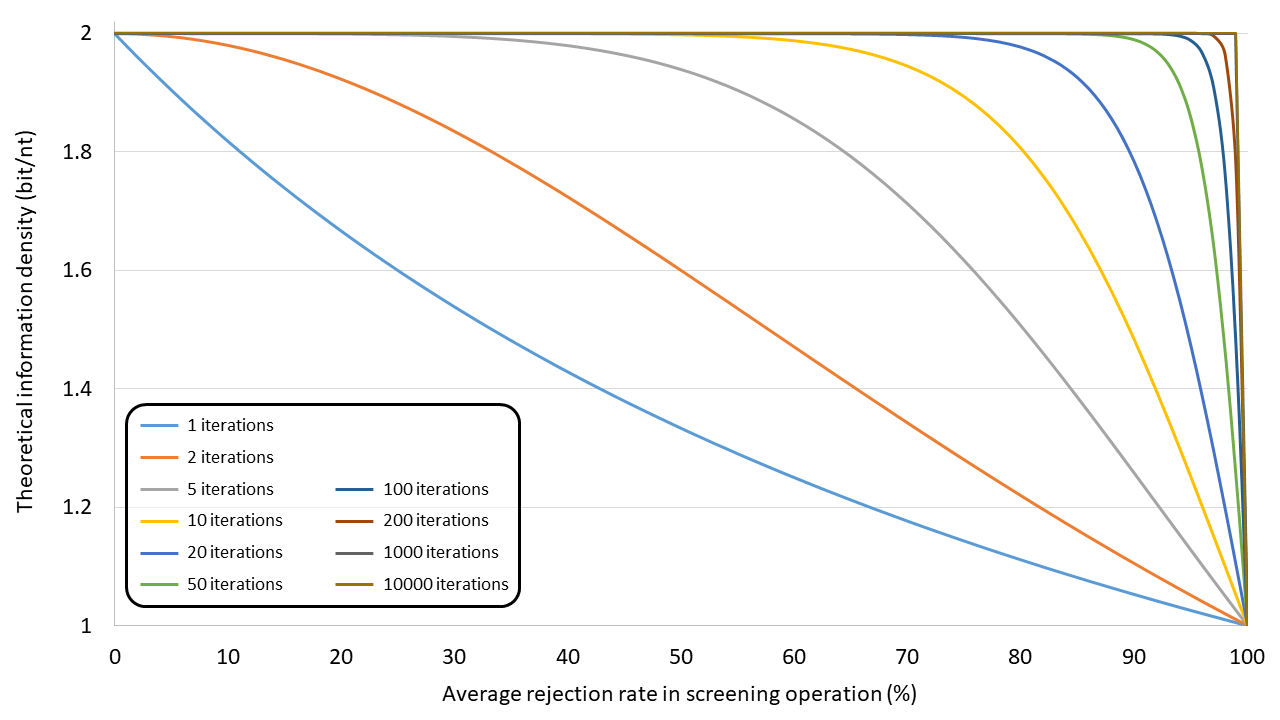

**Figure S2.** **The information density estimation and average rejection rate with different transcoding screening iterations.**

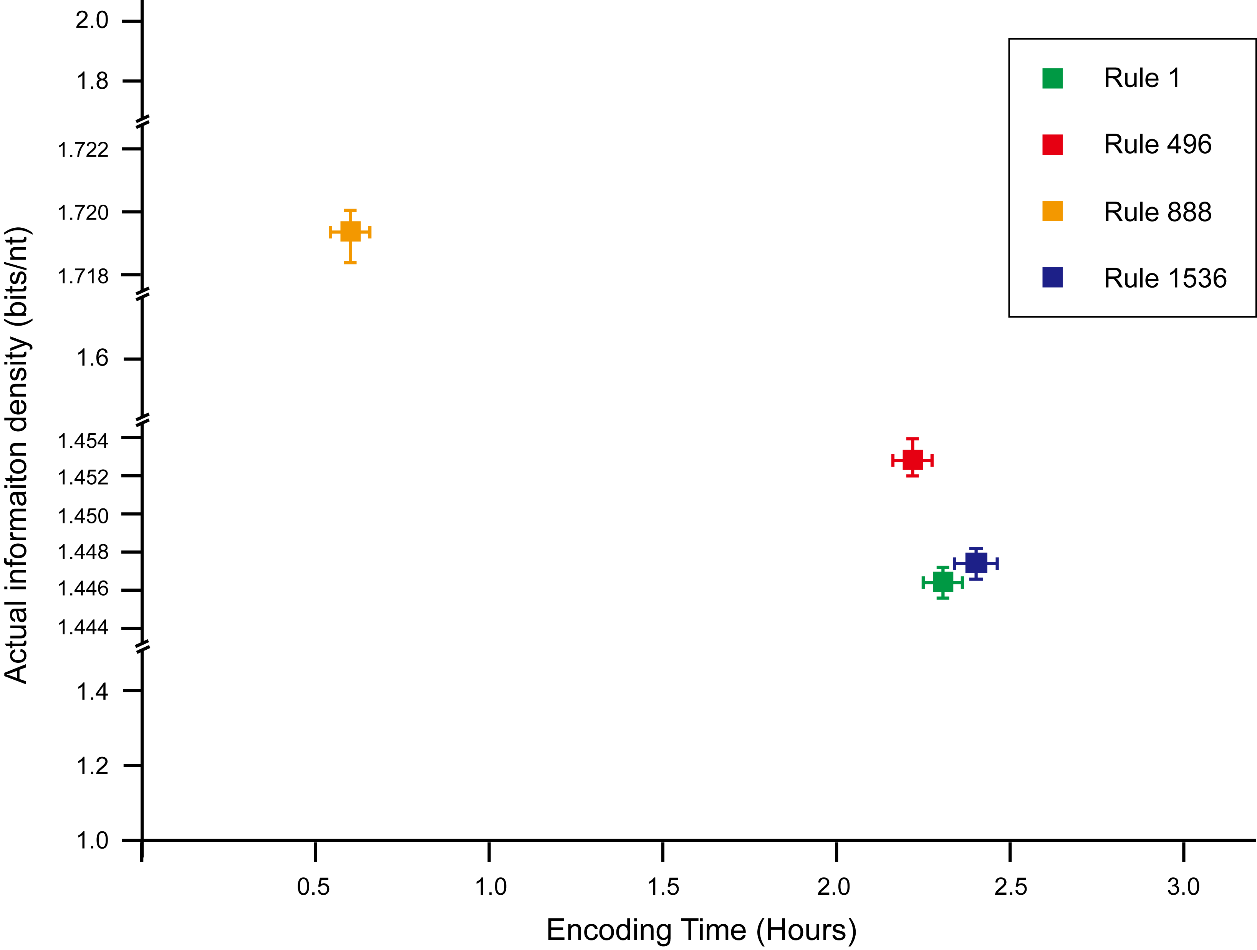

**Figure S3.** **The information density and transcoding time estimation of demo file ‘Exiting the Factory.flv’ using four randomly selected rules 1, 496, 888, and 1,536.**

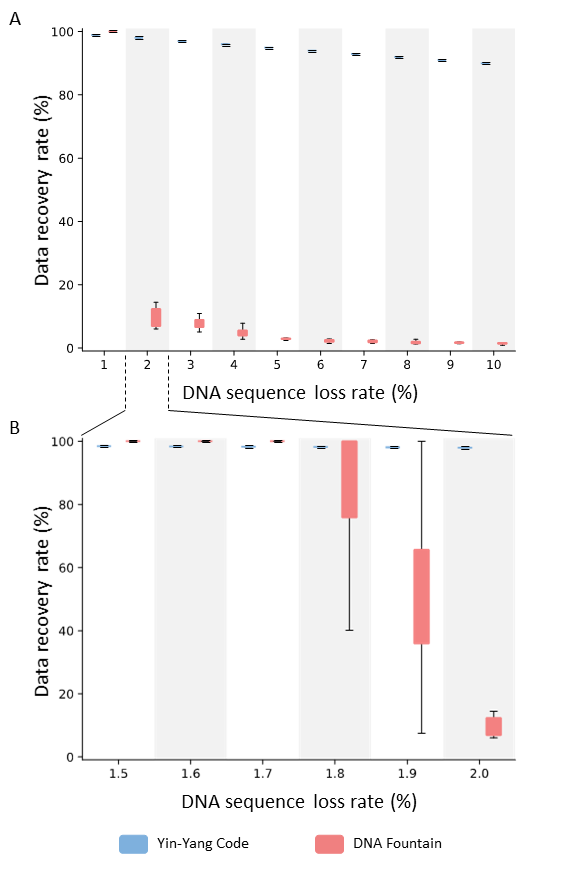

**Figure S4.** **The data recovery rate in the presence of a varying gradient of DNA sequence loss. The loss rate from A) 1% to 10% and B) 1.5% to 2.0%.**

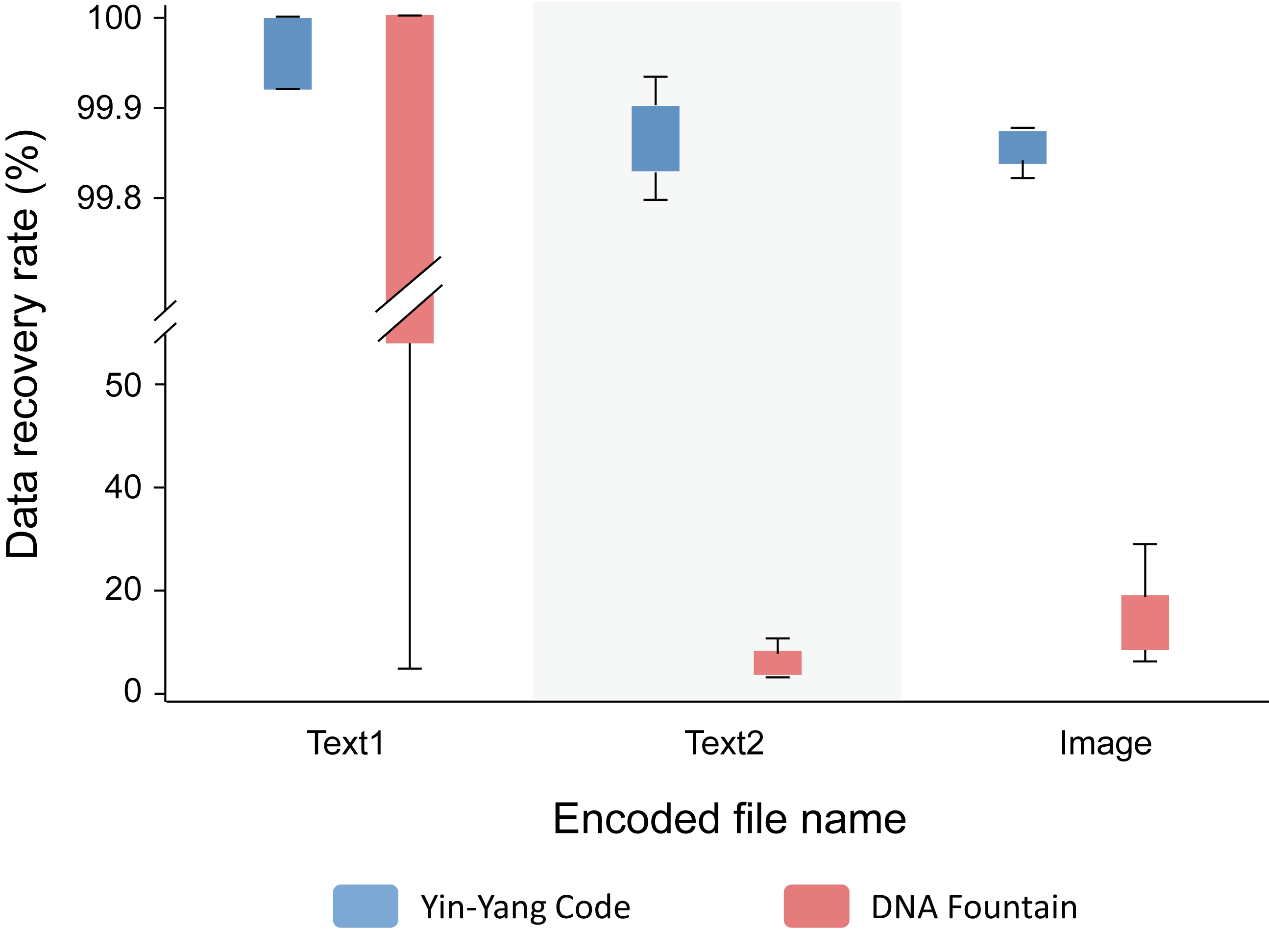

**Figure S5.** **The data recovery rate of three individual files according to experimental validation results with average copy number of 1,000.**

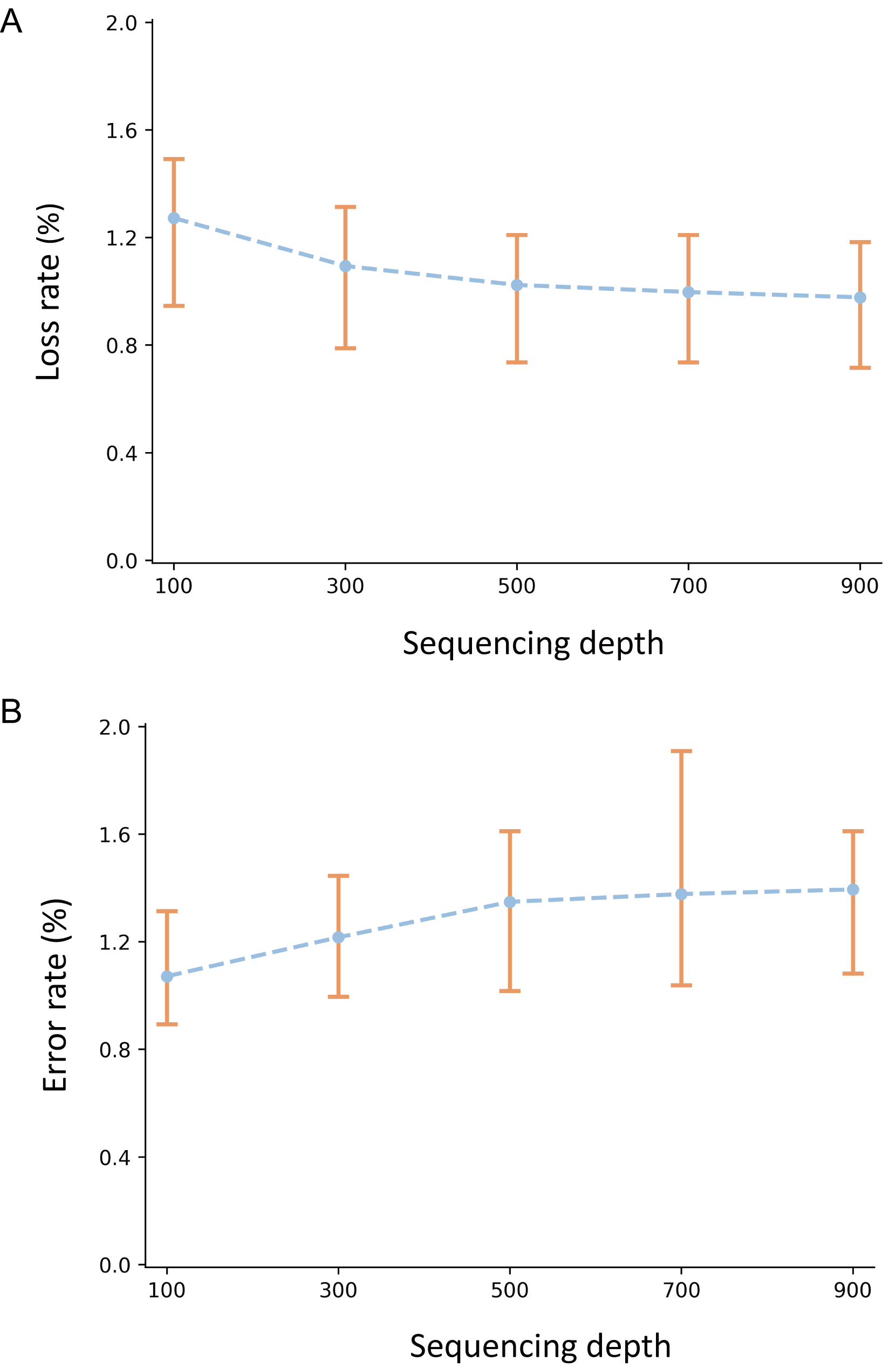

**Figure S6.** **Evaluation of the influence of sequencing depth on sequence error rate (A) and sequence loss rate (B) for in vitro storage validation.**

**Table S1.** ***The effect on compatibility of generated DNA sequences using different coding strategies for specific binary data patterns.***

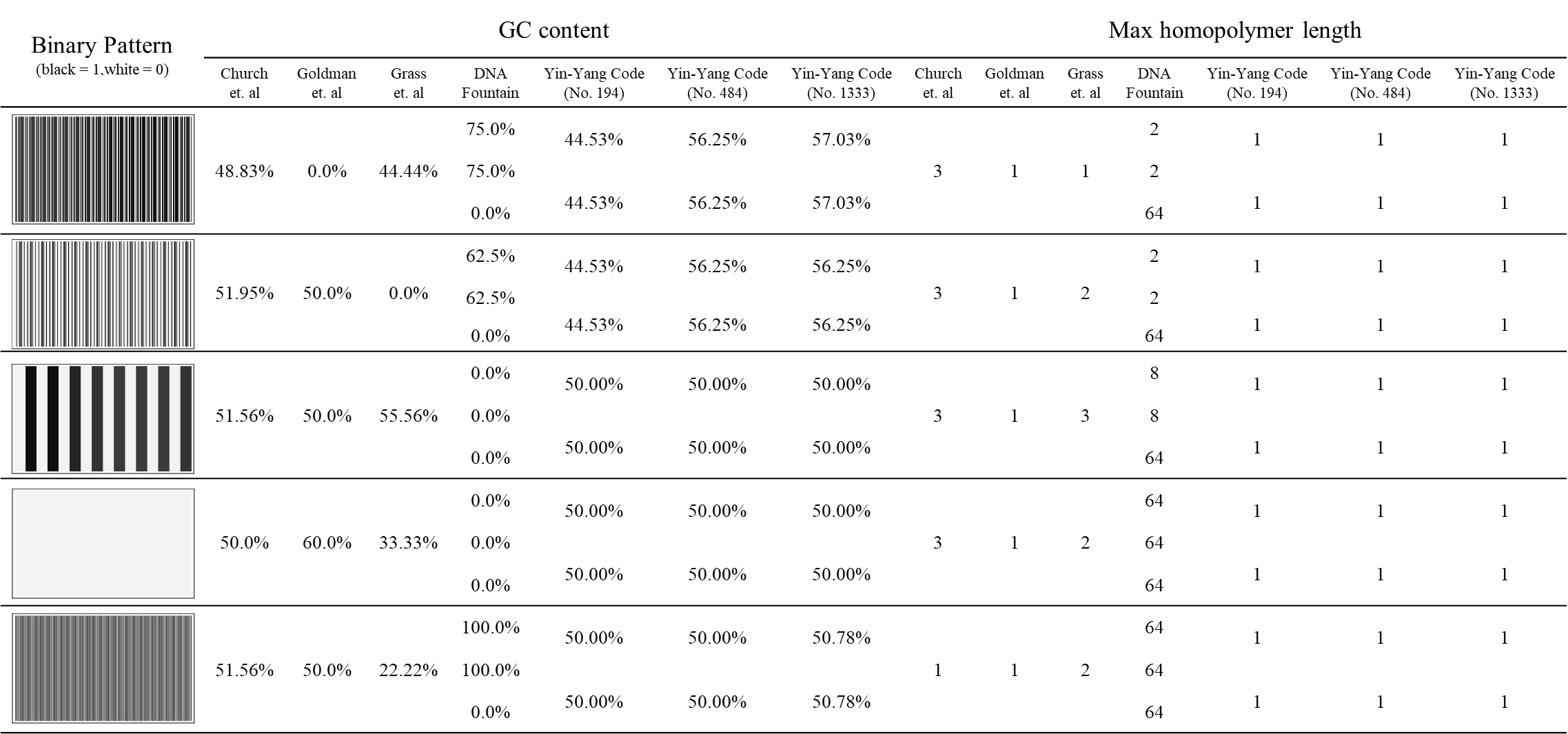

**Table S2.** **The estimation of iteration runs and corresponding information density by transcoding 10 different types/formats of files.**

| **File name** | **File format** | **File size (kB)** | **Rate of additional**  **information (%)** | **Average number of iteration runs** | **Information**  **density (bits/nt)**  **(Rule 496)** |
| --- | --- | --- | --- | --- | --- |
| Mona Lisa | jpg | 96 | 0.031 | 1.398 | 1.8037 |
| United Nations Flag | bmp | 469 | 0.075 | 5.040 | 1.7764 |
| A Tale of Two Cities | pdf | 1011 | 0.072 | 1.825 | 1.7506 |
| The Wandering Earth | pdf | 368 | 0.064 | 1.924 | 1.7766 |
| DNA Fountain Input Files | tar | 2096 | 0.004 | 1.493 | 1.7391 |
| Microsoft Winmine | exe | 117 | 3.180 | 7.873 | 1.7489 |
| For Elise | wma | 2544 | 0.462 | 3.482 | 1.7311 |
| Summer | mp3 | 6033 | 0.006 | 1.924 | 1.7265 |
| Exiting the Factory | flv | 4033 | 19.245 | 7.286 | 1.4480 |
| I have a Dream | mp4 | 3986 | 0.094 | 1.923 | 1.7250 |

**Table S3.** **The comparisons between YYC and DNA Fountain for transcoding 9 different types of files.** Minimum required redundancy for successful encoding and decoding larger than 100% is marked as “N.A.”.

| **File Names** | **Methods** | | | | |
| --- | --- | --- | --- | --- | --- |
|  | **Yin-Yang Code (Rule 496)** | | **DNA Fountain** | | |
|  | **Oligo Number** | **Net information Density (bits/nt)** | **Oligo Number** | **Minimum Redundancy** | **Net information Density (bits/nt)** |
| Mona Lisa | 3252 | 1.804 | 3811 | 25% | 1.347 |
| Microsoft Winmine | 4121 | 1.749 | 4980 | 33% | 1.266 |
| The Wandering Earth | 12536 | 1.777 | 13390 | 14% | 1.477 |
| United Nations Flag | 16014 | 1.776 | N.A. | | |
| A Tale of Two Cities | 34501 | 1.751 | 35554 | 10% | 1.531 |
| DNA fountain input files | 71513 | 1.739 | 71734 | 7% | 1.574 |
| For Elise | 87219 | 1.731 | 87090 | 7% | 1.574 |
| I Have a Dream | 136183 | 1.725 | 202808 | 59% | 1.059 |
| Summer | 205918 | 1.727 | N.A. | | |

**Table S4.** **Minimum redundancy required for successful data retrieval with different bitmap images.** Minimum required redundancy for successful encoding and decoding larger than 300% is marked as “N.A.”.

| **Name** | **Flag Style** | **Minimum Available Redundancy** |
| --- | --- | --- |
| United Nations Flag | 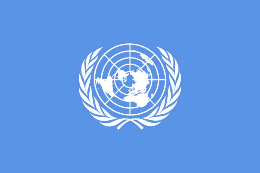 | > 66% |
| China Flag | 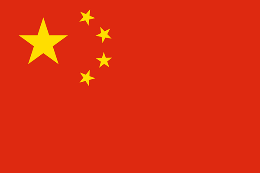 | > 243% |
| Japan Flag | 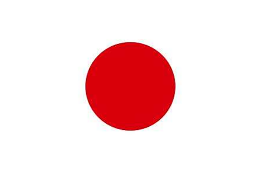 | N.A. |
| India Flag | 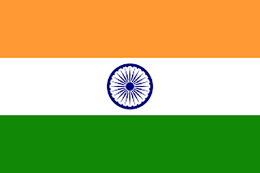 | > 42% |
| Britain Flag | 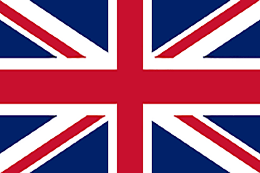 | 13% |
| Ireland Flag | 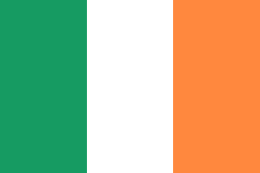 | N.A. |
| Germany Flag | 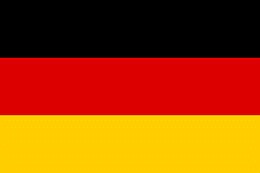 | N.A. |
| America Flag | 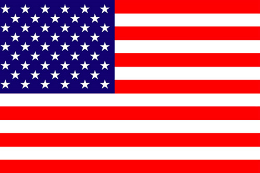 | > 61% |
| Brazil Flag | 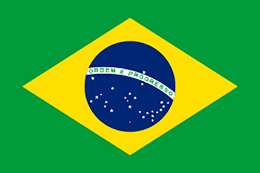 | > 116% |
| Singapore Flag | 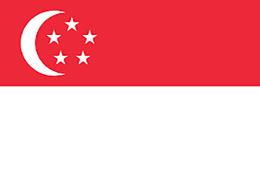 | 33% |

**Table S5.** **The data recovery statistics of YYC coding in vitro binary data storage validation.**

| **Encoded file** | **AMC number** | **Decoded file** | **Obtained**  **oligo amount** | **Required**  **oligo amount** | **Oligo coverage** | **Recovered binary**  **segments** | **Original binary**  **segments** | **Information recovery rate** |
| --- | --- | --- | --- | --- | --- | --- | --- | --- |
| Text1 | 10000 | YYC-File1-V300086833_L2_529.yyc_file1.dna | 1672 | 1676 | 99.76% | 2678 | 2678 | 100.00% |
| Text1 | 10000 | YYC-File1-V300086833_L2_530.yyc_file1.dna | 1672 | 1676 | 99.76% | 2678 | 2678 | 100.00% |
| Text1 | 10000 | YYC-File1-V300086833_L4_515.yyc_file1.dna | 1673 | 1676 | 99.82% | 2678 | 2678 | 100.00% |
| Text1 | 10000 | YYC-File1-V300086833_L4_501.yyc_file1.dna | 1670 | 1676 | 99.64% | 2676 | 2678 | 99.93% |
| Text1 | 10000 | YYC-File1-V300086833_L4_502.yyc_file1.dna | 1670 | 1676 | 99.64% | 2676 | 2678 | 99.93% |
| Text1 | 10000 | YYC-File1-V300086833_L4_516.yyc_file1.dna | 1672 | 1676 | 99.76% | 2676 | 2678 | 99.93% |
| Text1 | 1000 | YYC-File1-V300086833_L3_557.yyc_file1.dna | 1665 | 1676 | 99.34% | 2678 | 2678 | 100.00% |
| Text1 | 1000 | YYC-File1-V300086833_L3_558.yyc_file1.dna | 1664 | 1676 | 99.28% | 2678 | 2678 | 100.00% |
| Text1 | 1000 | YYC-File1-V300086833_L3_564.yyc_file1.dna | 1659 | 1676 | 98.99% | 2676 | 2678 | 99.93% |
| Text1 | 1000 | YYC-File1-V300086833_L3_563.yyc_file1.dna | 1658 | 1676 | 98.93% | 2676 | 2678 | 99.93% |
| Text1 | 100 | YYC-File1-V300086833_L3_578.yyc_file1.dna | 1281 | 1676 | 76.43% | 2164 | 2678 | 80.81% |
| Text1 | 100 | YYC-File1-V300086833_L3_577.yyc_file1.dna | 1271 | 1676 | 75.84% | 2135 | 2678 | 79.72% |
| Text1 | 100 | YYC-File1-V300086486_L3_583.yyc_file1.dna | 925 | 1676 | 55.19% | 1511 | 2678 | 56.42% |
| Text1 | 100 | YYC-File1-V300086486_L3_584.yyc_file1.dna | 916 | 1676 | 54.65% | 1494 | 2678 | 55.79% |
| Text1 | 100 | YYC-File1-V300086486_L4_584.yyc_file1.dna | 914 | 1676 | 54.53% | 1485 | 2678 | 55.45% |
| Text2 | 10000 | YYC-File2-V300086833_L4_518.yyc_file2.dna | 3796 | 3805 | 99.76% | 6083 | 6085 | 99.97% |
| Text2 | 10000 | YYC-File2-V300086833_L2_531.yyc_file2.dna | 3794 | 3805 | 99.71% | 6082 | 6085 | 99.95% |
| Text2 | 10000 | YYC-File2-V300086833_L2_532.yyc_file2.dna | 3795 | 3805 | 99.74% | 6082 | 6085 | 99.95% |
| Text2 | 10000 | YYC-File2-V300086833_L4_503.yyc_file2.dna | 3782 | 3805 | 99.40% | 6082 | 6085 | 99.95% |
| Text2 | 10000 | YYC-File2-V300086833_L4_504.yyc_file2.dna | 3783 | 3805 | 99.42% | 6081 | 6085 | 99.93% |
| Text2 | 10000 | YYC-File2-V300086833_L4_517.yyc_file2.dna | 3804 | 3805 | 99.97% | 6080 | 6085 | 99.92% |
| Text2 | 1000 | YYC-File2-V300086833_L3_560.yyc_file2.dna | 3767 | 3805 | 99.00% | 6080 | 6085 | 99.92% |
| Text2 | 1000 | YYC-File2-V300086833_L3_559.yyc_file2.dna | 3769 | 3805 | 99.05% | 6078 | 6085 | 99.88% |
| Text2 | 1000 | YYC-File2-V300086833_L3_566.yyc_file2.dna | 3752 | 3805 | 98.61% | 6074 | 6085 | 99.82% |
| Text2 | 1000 | YYC-File2-V300086833_L3_565.yyc_file2.dna | 3755 | 3805 | 98.69% | 6072 | 6085 | 99.79% |
| Text2 | 100 | YYC-File2-V300086833_L3_580.yyc_file2.dna | 3144 | 3805 | 82.63% | 5326 | 6085 | 87.53% |
| Text2 | 100 | YYC-File2-V300086833_L3_579.yyc_file2.dna | 3144 | 3805 | 82.63% | 5306 | 6085 | 87.20% |
| Text2 | 100 | YYC-File2-V300086486_L4_586.yyc_file2.dna | 2443 | 3805 | 64.20% | 4028 | 6085 | 66.20% |
| Text2 | 100 | YYC-File2-V300086486_L3_586.yyc_file2.dna | 2434 | 3805 | 63.97% | 4020 | 6085 | 66.06% |
| Text2 | 100 | YYC-File2-V300086486_L4_585.yyc_file2.dna | 2432 | 3805 | 63.92% | 4011 | 6085 | 65.92% |
| Text2 | 100 | YYC-File2-V300086486_L3_585.yyc_file2.dna | 2426 | 3805 | 63.76% | 3997 | 6085 | 65.69% |
| Image | 10000 | YYC-File3-V300086833_L4_519.yyc_file3.dna | 4607 | 4622 | 99.68% | 7275 | 7279 | 99.95% |
| Image | 10000 | YYC-File3-V300086833_L4_520.yyc_file3.dna | 4603 | 4622 | 99.59% | 7275 | 7279 | 99.95% |
| Image | 10000 | YYC-File3-V300086833_L2_542.yyc_file3.dna | 4601 | 4622 | 99.55% | 7273 | 7279 | 99.92% |
| Image | 10000 | YYC-File3-V300086833_L2_541.yyc_file3.dna | 4607 | 4622 | 99.68% | 7271 | 7279 | 99.89% |
| Image | 10000 | YYC-File3-V300086833_L4_505.yyc_file3.dna | 4596 | 4622 | 99.44% | 7270 | 7279 | 99.88% |
| Image | 10000 | YYC-File3-V300086833_L4_506.yyc_file3.dna | 4595 | 4622 | 99.42% | 7270 | 7279 | 99.88% |
| Image | 1000 | YYC-File3-V300086833_L3_567.yyc_file3.dna | 4569 | 4622 | 98.85% | 7268 | 7279 | 99.85% |
| Image | 1000 | YYC-File3-V300086833_L3_561.yyc_file3.dna | 4569 | 4622 | 98.85% | 7268 | 7279 | 99.85% |
| Image | 1000 | YYC-File3-V300086833_L3_568.yyc_file3.dna | 4570 | 4622 | 98.87% | 7267 | 7279 | 99.84% |
| Image | 1000 | YYC-File3-V300086833_L3_562.yyc_file3.dna | 4578 | 4622 | 99.05% | 7266 | 7279 | 99.82% |
| Image | 100 | YYC-File3-V300086486_L4_581.yyc_file3.dna | 3670 | 4622 | 79.40% | 6137 | 7279 | 84.31% |
| Image | 100 | YYC-File3-V300086486_L3_581.yyc_file3.dna | 3659 | 4622 | 79.16% | 6131 | 7279 | 84.23% |
| Image | 100 | YYC-File3-V300086486_L4_582.yyc_file3.dna | 3653 | 4622 | 79.04% | 6110 | 7279 | 83.94% |
| Image | 100 | YYC-File3-V300086486_L3_582.yyc_file3.dna | 3653 | 4622 | 79.04% | 6109 | 7279 | 83.93% |
| Image | 100 | YYC-File3-V300086486_L4_588.yyc_file3.dna | 2789 | 4622 | 60.34% | 4560 | 7279 | 62.65% |
| Image | 100 | YYC-File3-V300086486_L3_588.yyc_file3.dna | 2791 | 4622 | 60.39% | 4554 | 7279 | 62.56% |
| Image | 100 | YYC-File3-V300086486_L3_587.yyc_file3.dna | 2755 | 4622 | 59.61% | 4504 | 7279 | 61.88% |
| Image | 100 | YYC-File3-V300086486_L4_587.yyc_file3.dna | 2750 | 4622 | 59.50% | 4496 | 7279 | 61.77% |

**Table S6.** **The data recovery statistics of DNA Fountain coding in vitro binary data storage validation.**

| **Encoded file** | **AMC number** | **Decoded file** | **Obtained**  **oligo amount** | **Required**  **oligo amount** | **Oligo coverage** | **Recovered binary**  **segments** | **Original binary**  **segments** | **Information recovery rate** |
| --- | --- | --- | --- | --- | --- | --- | --- | --- |
| Text1 | 10000 | DF-S-File1-WS2-0-V300086833_L2_543.df-s-file1.dna | 1634 | 1634 | 100.00% | 1339 | 1339 | 100.00% |
| Text1 | 10000 | DF-S-File1-WS2-0-V300086833_L2_544.df-s-file1.dna | 1634 | 1634 | 100.00% | 1339 | 1339 | 100.00% |
| Text1 | 10000 | DF-S-File1-WS2-0-V300086833_L4_507.df-s-file1.dna | 1634 | 1634 | 100.00% | 1339 | 1339 | 100.00% |
| Text1 | 10000 | DF-S-File1-WS2-0-V300086833_L4_508.df-s-file1.dna | 1634 | 1634 | 100.00% | 1339 | 1339 | 100.00% |
| Text1 | 10000 | DF-S-File1-WS2-0-V300086833_L4_521.df-s-file1.dna | 1634 | 1634 | 100.00% | 1339 | 1339 | 100.00% |
| Text1 | 10000 | DF-S-File1-WS2-0-V300086833_L4_522.df-s-file1.dna | 1634 | 1634 | 100.00% | 1339 | 1339 | 100.00% |
| Text1 | 1000 | DF-S-File1-WS2-1-20201229-2.df-s_f1.dna | 1634 | 1634 | 100.00% | 1339 | 1339 | 100.00% |
| Text1 | 1000 | DF-S-File1-WS2-1-20201229-3.df-s_f1.dna | 1634 | 1634 | 100.00% | 1339 | 1339 | 100.00% |
| Text1 | 1000 | DF-S-File1-WS2-1-20201229-1.df-s_f1.dna | 1630 | 1634 | 99.76% | 43 | 1339 | 3.21% |
| Text1 | 100 | DF-S-File1-WS2-2-20201229-2.df-s_f1.dna | 834 | 1634 | 51.04% | 10 | 1339 | 0.75% |
| Text1 | 100 | DF-S-File1-WS2-2-20201229-3.df-s_f1.dna | 681 | 1634 | 41.68% | 15 | 1339 | 1.12% |
| Text1 | 100 | DF-S-File1-WS2-2-20201229-1.df-s_f1.dna | 389 | 1634 | 23.81% | 8 | 1339 | 0.60% |
| Text2 | 10000 | DF-S-File2-WS2-0-V300086833_L2_546.df-s-file2.dna | 5262 | 5265 | 99.94% | 3040 | 3040 | 100.00% |
| Text2 | 10000 | DF-S-File2-WS2-0-V300086833_L4_509.df-s-file2.dna | 5262 | 5265 | 99.94% | 3040 | 3040 | 100.00% |
| Text2 | 10000 | DF-S-File2-WS2-0-V300086833_L2_523.df-s-file2.dna | 5262 | 5265 | 99.94% | 3040 | 3040 | 100.00% |
| Text2 | 10000 | DF-S-File2-WS2-0-V300086833_L2_545.df-s-file2.dna | 5261 | 5265 | 99.92% | 3040 | 3040 | 100.00% |
| Text2 | 10000 | DF-S-File2-WS2-0-V300086833_L4_510.df-s-file2.dna | 5260 | 5265 | 99.91% | 3040 | 3040 | 100.00% |
| Text2 | 10000 | DF-S-File2-WS2-0-V300086833_L2_524.df-s-file2.dna | 5258 | 5265 | 99.87% | 3040 | 3040 | 100.00% |
| Text2 | 1000 | DF-S-File2-WS2-1-20201229-2.df-s_f2.dna | 5246 | 5265 | 99.64% | 155 | 3040 | 5.10% |
| Text2 | 1000 | DF-S-File2-WS2-1-20201229-1.df-s_f2.dna | 5227 | 5265 | 99.28% | 15 | 3040 | 0.49% |
| Text2 | 1000 | DF-S-File2-WS2-1-20201229-3.df-s_f2.dna | 5223 | 5265 | 99.20% | 17 | 3040 | 0.56% |
| Text2 | 100 | DF-S-File2-WS2-2-20201229-2.df-s_f2.dna | 2426 | 5265 | 46.08% | 4 | 3040 | 0.13% |
| Text2 | 100 | DF-S-File2-WS2-2-20201229-3.df-s_f2.dna | 1663 | 5265 | 31.59% | 3 | 3040 | 0.10% |
| Text2 | 100 | DF-S-File2-WS2-2-20201229-1.df-s_f2.dna | 685 | 5265 | 13.01% | 4 | 3040 | 0.13% |
| Image | 10000 | DF-S-File3-WS2-0-V300086833_L4_511.df-s-file3.dna | 4076 | 4077 | 99.98% | 3640 | 3640 | 100.00% |
| Image | 10000 | DF-S-File3-WS2-0-V300086833_L2_547.df-s-file3.dna | 4074 | 4077 | 99.93% | 3640 | 3640 | 100.00% |
| Image | 10000 | DF-S-File3-WS2-0-V300086833_L2_525.df-s-file3.dna | 4073 | 4077 | 99.90% | 3640 | 3640 | 100.00% |
| Image | 10000 | DF-S-File3-WS2-0-V300086833_L2_526.df-s-file3.dna | 4073 | 4077 | 99.90% | 3640 | 3640 | 100.00% |
| Image | 10000 | DF-S-File3-WS2-0-V300086833_L4_512.df-s-file3.dna | 4073 | 4077 | 99.90% | 3640 | 3640 | 100.00% |
| Image | 10000 | DF-S-File3-WS2-0-V300086833_L2_548.df-s-file3.dna | 4072 | 4077 | 99.88% | 3640 | 3640 | 100.00% |
| Image | 1000 | DF-S-File3-WS2-1-20201229-3.df-s_f3.dna | 4071 | 4077 | 99.85% | 81 | 3640 | 2.23% |
| Image | 1000 | DF-S-File3-WS2-1-20201229-2.df-s_f3.dna | 4069 | 4077 | 99.80% | 802 | 3640 | 22.03% |
| Image | 1000 | DF-S-File3-WS2-1-20201229-1.df-s_f3.dna | 4066 | 4077 | 99.73% | 169 | 3640 | 4.64% |
| Image | 100 | DF-S-File3-WS2-2-20201229-2.df-s_f3.dna | 2119 | 4077 | 51.97% | 11 | 3640 | 0.30% |
| Image | 100 | DF-S-File3-WS2-2-20201229-3.df-s_f3.dna | 1650 | 4077 | 40.47% | 7 | 3640 | 0.19% |
| Image | 100 | DF-S-File3-WS2-2-20201229-1.df-s_f3.dna | 1335 | 4077 | 32.74% | 13 | 3640 | 0.36% |
| Tar file | 10000 | DF-tar-WS3-1-V300086833_L2_550.df-tar.dna | 9057 | 9185 | 98.61% | 8129 | 8129 | 100.00% |
| Tar file | 10000 | DF-tar-WS3-1V300086833_L2_549.df-tar.dna | 9052 | 9185 | 98.55% | 8129 | 8129 | 100.00% |
| Tar file | 10000 | DF-tar-WS3-1V300086833_L4_513.df-tar.dna | 8953 | 9185 | 97.47% | 8129 | 8129 | 100.00% |
| Tar file | 10000 | DF-tar-WS3-1V300086833_L4_514.df-tar.dna | 8861 | 9185 | 96.47% | 8129 | 8129 | 100.00% |
| Tar file | 1000 | DF-tar-WS3-1-20201229-2.df-tar.dna | 9048 | 9185 | 98.51% | 2669 | 8129 | 32.83% |
| Tar file | 1000 | DF-tar-WS3-1-20201229-3.df-tar.dna | 9021 | 9185 | 98.21% | 57 | 8129 | 0.70% |
| Tar file | 1000 | DF-tar-WS3-1-20201229-1.df-tar.dna | 9016 | 9185 | 98.16% | 215 | 8129 | 2.64% |
| Tar file | 100 | DF-tar-WS3-2-20201229-2.df-tar.dna | 5209 | 9185 | 56.71% | 5 | 8129 | 0.06% |
| Tar file | 100 | DF-tar-WS3-2-20201229-3.df-tar.dna | 3616 | 9185 | 39.37% | 11 | 8129 | 0.14% |
| Tar file | 100 | DF-tar-WS3-2-20201229-1.df-tar.dna | 1068 | 9185 | 11.63% | 4 | 8129 | 0.05% |

**Table S7.** **Error analysis of in vivo storage demonstration. Sequences are mapping to original sequence. SNP refers to substitution of a single nucleotide. Indel refers to insertion or deletion of a single nucleotide. Structural variation refers to the variation in structure; in this work, only deletion was observed.**

| Strain ID | Single nucleotide polymorphism (SNP) | 1-nt Indel | Structural Variation (SV) |
| --- | --- | --- | --- |
| L01 | 140 | 26 | 4 |
| L02 | 146 | 41 | 2 |
| L03 | 122 | 37 | 1 |

**Command line steps to encode and decode data**

For encoding process

\# set pre-redundancy in digital data

redundancy_factor = 4

binary_segments_with_redundancy = []

for i in range(0, len(binary_segments), redundancy_factor):

binary_segments_with_redundancy += binary_segments[i:i+redundancy_factor] + \

[self.xor_segments(binary_segments[i:i+redundancy_factor])]

\# set rule 888

[support_base, rule1, rule2] = ["A", [1, 0, 0, 1], [[0, 1, 0, 1], [1, 1, 0, 0], [1, 0, 1, 0], [0, 0, 1, 1]]]

\# initialize Yin-Yang Code

tool=yyc.YYC(support_bases=support_base,base_reference=rule1, current_code_matrix=rule2,

max_homopolymer=4,max_content=0.6,min_free_energy=-30,

search_count=100)

\# encoding by codec factory

codec_factory.encode(method=tool,input_path=read_file_path,output_path=dna_path,

model_path=model_path,need_index=True,need_log=True)

For decoding process

\# initialize Yin-Yang Code

tool=yyc.YYC(support_bases=support_base,base_reference=rule1, current_code_matrix=rule2,

max_homopolymer=4,max_content=0.6,min_free_energy=-30,

search_count=100)

\# decoding by codec factory

codec_factory.decode(method=tool,input_path=dna_path,output_path=write_file_path,

has_index=True,need_log=True)

For verification process

\# reading source and target digital data

matrix_1, _ = data_handle.read_binary_from_all(read_file_path, 120, False)

matrix_2, _ = data_handle.read_binary_from_all(write_file_path, 120, False)

\# comparison

print("source digital file == target digital file: " + str(matrix_1 == matrix_2))
